# Machine learning of stochastic gene network phenotypes

**DOI:** 10.1101/825943

**Authors:** Kyemyung Park, Thorsten Prüstel, Yong Lu, John S. Tsang

**Author notes:** To whom correspondence should be addressed (JST).

## Abstract

A recurrent challenge in biology is the development of predictive quantitative models because most molecular and cellular parameters have unknown values and realistic models are analytically intractable. While the dynamics of the system can be analyzed via computer simulations, substantial computational resources are often required given uncertain parameter values resulting in large numbers of parameter combinations, especially when realistic biological features are included. Simulation alone also often does not yield the kinds of intuitive insights from analytical solutions. Here we introduce a general framework combining stochastic/mechanistic simulation of reaction systems and machine learning of the simulation data to generate computationally efficient predictive models and interpretable parameter-phenotype maps. We applied our approach to investigate stochastic gene expression propagation in biological networks, which is a contemporary challenge in the quantitative modeling of single-cell heterogeneity. We found that accurate, predictive machine-learning models of stochastic simulation results can be constructed. Even in the simplest networks existing analytical schemes generated significantly less accurate predictions than our approach, which revealed interesting insights when applied to more complex circuits, including the extensive tunability of information propagation enabled by feedforward circuits and how even single negative feedbacks can utilize stochastic fluctuations to generate robust oscillations. Our approach is applicable beyond biology and opens up a new avenue for exploring complex dynamical systems.

---

A major goal of systems biology is to develop quantitative models to predict the behavior of biological systems (*1, 2*). However, most realistic molecular and cellular models have a large number of parameters (e.g., reaction rates, cellular proliferation rates, extent of physical interactions among cells or molecules), whose values remain unknown and are often challenging to measure or infer quantitatively (*3, 4*). While some biological phenotypes are robust to parameter variations (*5*), most are “tunable” by parameters (*6*). Therefore, analyzing the behavior of a system over the *entire* plausible space of parameters is needed to study the phenotypic range of a biological system and its parameter-phenotype relationships (*7–9*). A case in point concerns a contemporary problem in single cell biology: Despite the increasing availability of single-cell gene expression data enabled by rapid technological advances (*10*), an important unanswered question is how cell-to-cell expression variation and gene-gene correlation among single cells are regulated by the underlying gene regulatory network (GRN), within which different signals, including those arising from environmental variations or biochemical fluctuations, are transmitted (*11–13*).

Stochastic master equations (SMEs) (see Glossary) can be used to model, analyze, and predict single-cell heterogeneity and gene-gene correlation behaviors based on GRNs. A multitude of cell type- and environment-dependent stoichiometric and kinetic parameters are required, but their values remain largely unknown. To make SMEs analytically tractable, simplifying assumptions are needed with the risk of ignoring important features such as “bursty” transcription (*14, 15*). Alternatively, resource intensive computational simulations (e.g., using Gillespie’s Algorithm (*16*)) can be used to obtain the stochastic dynamics of individual parameter configurations; sensitivity analysis can then be employed to evaluate which parameters can affect the phenotype of interest (*3*). However, extensive computational resources are required to analyze the parameter space, given the large uncertainty in parameter values and the complex correlation structure among parameters. Plus, simulation analysis alone often does not automatically yield intuitive understanding.

Here, we combine computational simulation of full-feature stochastic models and machine learning (ML) to develop a framework, called MAchine learning of Parameter Phenotype Analysis (MAPPA), for constructing, exploring, and analyzing the mapping between parameters and quantitative phenotypes of a stochastic dynamical system (Figures 1A and S1; Supplementary text). Our goal is to take advantage of the large amounts of data that can be generated from bottom-up, mechanistic computational simulation of dynamical systems and the ability of modern machine learning approaches to “compress” such data to generate computationally efficient and interpretable models. MAPPA thus builds efficient, predictive, and interpretable ML models that capture the nonlinear mapping between parameter and phenotypic spaces (parameter-phenotype maps). The ML models can be viewed as “phenomenological” solutions of the SME that can predict the system’s quantitative behavior from parameter combinations, thus bypassing computationally expensive simulations. They also can delineate which and how parameters and parameter combinations shape phenotypes, both globally throughout the parameter space and locally in the neighborhoods of individual parameter configurations. We introduce visualizations to enable interactive exploration of the parameter-phenotype map, including a web application for interrogations of analysis below (https://phasespace-explorer.niaid.nih.gov).

**Figure 1.**
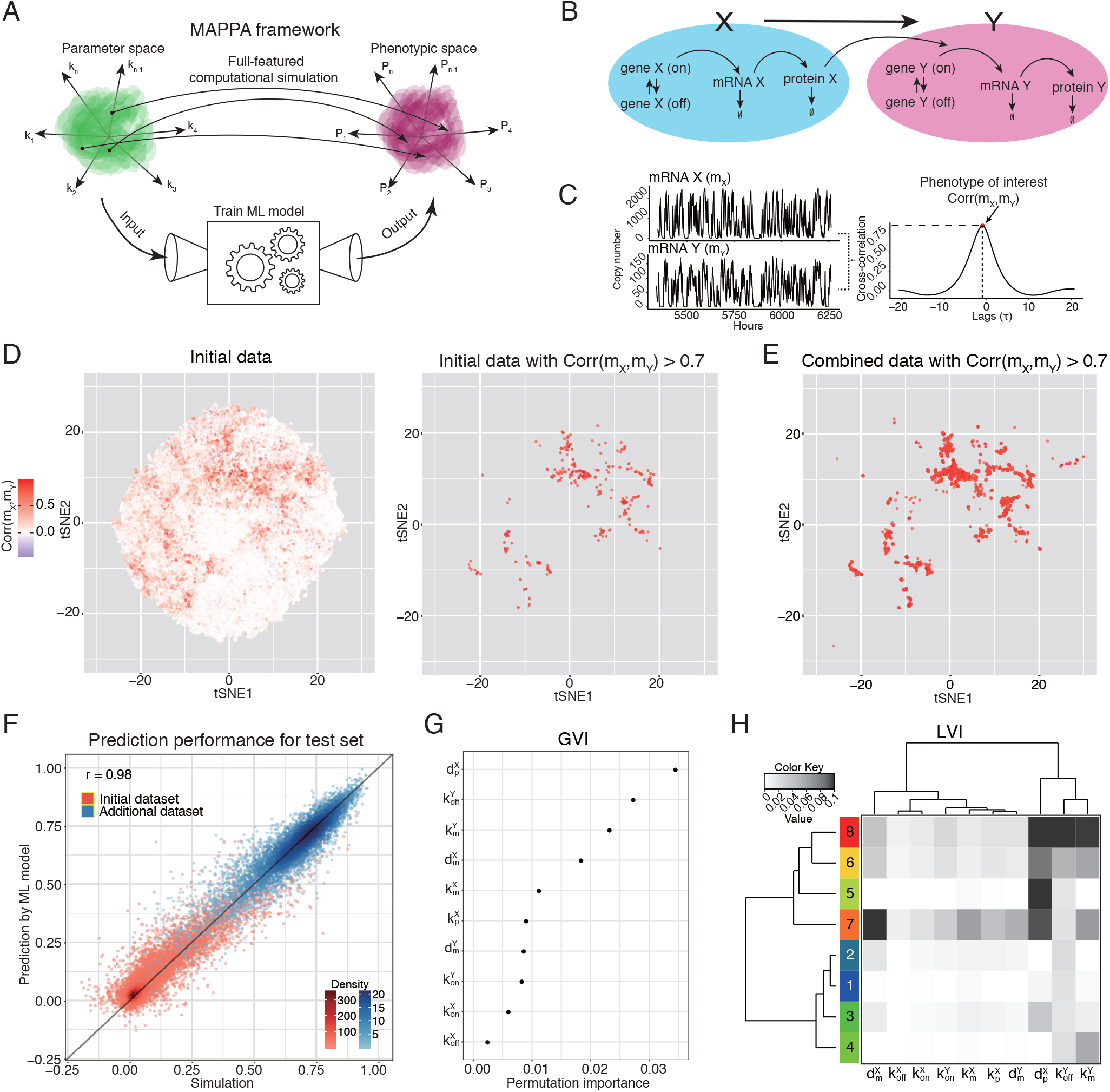
MAPPA overview and applying it to study the propagation of gene expression variability in a two-gene (transcription factor-target gene) network. **(A)** MAPPA utilizes massive simulation data and machine learning to construct models that can accurately and efficiently predict quantitative phenotypes given high-dimensional parameter combinations without using resource intensive dynamical simulations. The resulting machine learning models also serve as interpretable parameter-phenotype maps. **(B)** The two-gene network model in which the protein product of gene X regulates the transcription rate of gene Y. The promoter of both X and Y undergoes stochastic on-off state switching. Both the mRNAs and proteins undergo first order degradation. See Methods for additional details. **(C)** Definition of the phenotype of interest: quantifying the propagation of gene expression variability (or information) from gene X to gene Y. The time trajectories of mRNA X and Y generated by stochastic simulation from a specific parameter configuration is shown here for illustration. Here the metric of information transmission/propagation used, *Corr*(*m_X_, m_Y_*), was defined as the maximum of the cross-correlation between *m_X_*(*t*) (level of the mRNA X) and *m_Y_*(*t – τ*) (level of the mRNA Y) across a pre-defined range of time lags *τ*, (here the red dot indicates the maximum across, *τ*s). We only consider correlations with, *τ* 0 in this network to capture the causal relationship between X and Y (X ➔ Y). **(D and E)** Visualization of the parameter space and the phenotype using two-dimensional (2d) t-distributed Stochastic Neighbor Embedding (tSNE) computed from the sampled parameter combinations. (D) tSNE plot for (left) the initial simulated data and its subset (right) with *Corr*(*m_X_, m_Y_*) > 0.7. (E) Additional parameter combinations nearby those with *Corr*(*m_X_, m_Y_*) > 0.7 were sampled (referred herein as “additionally sampled”) and simulated to increase the representation of high-correlation parameter combinations; here the tSNE plot for the combined data (initial samples plus additionally sampled) with *Corr*(*m_X_, m_Y_*) > 0.7 is shown. The color scale denotes *Corr*(*m_X_, m_Y_*). **(F)** Scatter plot showing the concordance between the independently simulated *Corr*(*m_X_, m_Y_*) (x axis – not used in ML model training) vs. those predicted by the RF regression model using the model parameter values alone (y axis) (r = 0.98); note that the RF model was trained using the combined training data; each point corresponds to a parameter combination. The red points are those from the independent test set of the initially sampled parameter combinations; the blue dots correspond to the independent test set of the additionally sampled parameter combinations nearby the initial combinations with high (>0.7) *Corr*(*m_X_,m_Y_*). The color scale denotes the distribution density reflecting the relative abundance of data points (see Methods for details). **(G)** Global Variable Importance (GVI) (x axis) of the model parameters (y axis) fitted by the RF regression model. The permutation GVI is shown, which reflects the increase in prediction errors in out-of-bag data after permuting the indicated variable. Another type of GVI (impurity GVI) is shown together with the permutation GVI in Figure S2G. **(H)** A summary heatmap depicting the Local Variable Importance (LVI) of the RF regression model. Each row corresponds to a cluster of parameter combinations that exhibit similar LVI profiles across the indicated parameters (columns); the values shown in the heatmap are the average across all parameter combinations within each cluster. Eight clusters are shown as indicated by the cluster number/color bar on the left; the number of clusters were chosen qualitatively by considering: 1) ease of visualization in the limited space, 2) the qualitative diversity of LVIs that the clusters can illustrate (Methods). The LVI values shown in the heatmap are the average increases in the squared out-of-bag residuals provided by the *randomForest* package in R (see Methods).

To assess MAPPA, we first applied it to study how information (as encoded by the changes in gene expression over time) is transmitted from one gene to another (i.e., the propagation of variation (PoV)) in a prototypical, two-gene network model in single cells (Figure 1B, see Methods) (*12*). While existing analytically tractable models for this circuit require simplifying assumptions (*14, 15*), here we analyzed a full-feature model with mRNAs and proteins as distinct species, and promoters that can be switched stochastically between transcriptionally active (on) and inactive (off) states. As a measure of information propagation between genes X and Y, we defined the maximum time-lagged correlation (denoted as *Corr*(*m_X_*, *m_Y_*)) between mRNAs X and Y as the maximum cross-correlation between *m_X_*(*t*) and *m_Y_*(*t – τ*), where *m_X_* (*t*) and *x_Y_*(*t*) are copy numbers of mRNAs X and Y at time *t*, respectively, and *τ*, is the time lag (Figure 1C). The same metric can be applied to proteins, here we chose to focus on mRNAs since they are the dominant measurement modality in single cell studies.

Simulation on a large number (76,532) of randomly sampled, biologically plausible parameter combinations (see Methods) revealed that very few had high correlations (e.g., only 315 had a correlation of greater than 0.7) (Figure S2A) (*12*). Dimension reduction visualization using tSNE (Figure 1D) indicated that the parameters with high *Corr*(*m_X_, m_Y_*) formed clusters. Moreover, additional parameter combinations sampled from these regions also had high correlations. Thus, biased sampling guided by the phenotype of the neighbors can be used to increase the representation of rare parameter combinations (Figure 1E and S2B). Using this approach, we trained random forest (RF) regression models for *Corr*(*m_X_, m_Y_*) and assessed their predictive capacity using independently simulated data (Figure S2C). Both the model trained using the initial, uniformly sampled parameter combinations (r=0.93; Figure S2D) and the one trained by incorporating additional samples from the high-correlation regions showed excellent prediction performance (r = 0.98; Figure 1F); the latter had better performance in the high-correlation regions (Figure S2E). A two-class (high versus low correlations), categorical RF model performed similarly well (AUC = 0.98; Figure S2F; see Methods).

RF ML models provide “variable importance” to quantify the extent of influence a parameter can exert on the phenotype, both globally (GVI, Figures 1G and S2G) for the entire parameter space and locally (LVI, Figures 1H and S2H) at a particular point in that space (see Methods). For example, GVI revealed that the degradation rate of protein X 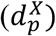, the off-rate of the promoter of gene Y 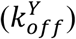, and the transcription rate of mRNA Y 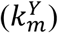 are the most important for determining *Corr*(*m_X_, m_Y_*), while the promoter switching rates of gene X (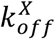 and 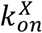) are less important (Figure 1G). Based on the LVI profiles, parameter combinations can be clustered into qualitatively distinct groups (Figures 1H and S2H). For example, the degradation rate of mRNA X is more important in cluster 7 than in other clusters (Figure 1H). Thus, individual parameters can exert local, “neighborhood”-dependent influences on the phenotype, consistent with the notion that gene networks may employ distinct strategies for PoV regulation depending on the cellular and environmental conditions (*12*).

To assess MAPPA further, we examined an analytical model (*14, 17*) in which simplifying assumptions were made to attain tractability (e.g., the promoter on/off switching is averaged; see Methods.) This model showed that 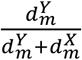 is one of the main factors determining *Corr*(*m_X_,m_Y_*) (see Methods), which is consistent with the GVI that 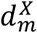 is important. However, LVI predicted that the effect of 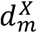 on *Corr*(*m_X_,m_Y_*) depends on the actual parameter combinations (Figure 1H). Notably, it is important in LVI cluster 7 but less so in cluster 8. In both clusters (more pronounced in cluster 8), the ML model performed better than the analytical model (Figures 2A and 2B). The analytical model had incorrectly predicted that those parameter combinations with 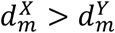 (prevalent in cluster 8) have low *Corr*(*m_X_,m_Y_*) (Figures 2B for cluster 8 and S3A-B for all parameter combinations). Closer examination revealed that these parameter configurations also had high 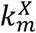 and 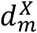 relative to the promoter switching rates (i.e., 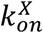 and 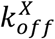). Thus, the promotor switching dynamics of gene X was a main driver of the fluctuations of mRNA X: a burst of transcripts was made in the on-state and transcripts were then degraded rapidly in the off-state. This gave rise to non-Poissonian mRNA fluctuations (*18*) and coordinated, “discrete” states involving two genes (i.e., either low or high levels of both mRNAs X and Y), as exemplified in the case shown in Figures 2C and 2D (parameter key: 042015_AAACEZGP; see Methods for more information on parameter keys). In contrast, those parameter combinations with 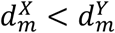 exhibited more continuous correlations (see Figures S3C-D for an example). This PoV mechanism may underlie the discrete single-cell gene-gene correlations observed in our earlier experiments (*12*) and could be employed by cells to attain multi-stable gene expression states even without complex feedback/feedforward mechanisms (*19*). “Bursty” transcriptional dynamics is a well-known source of cell-to-cell expression variation of individual genes, here MAPPA analysis revealed fresh insights on how such variations can be propagated from gene to gene in the regulatory network to generate distinct cellular states.

**Figure 2.**
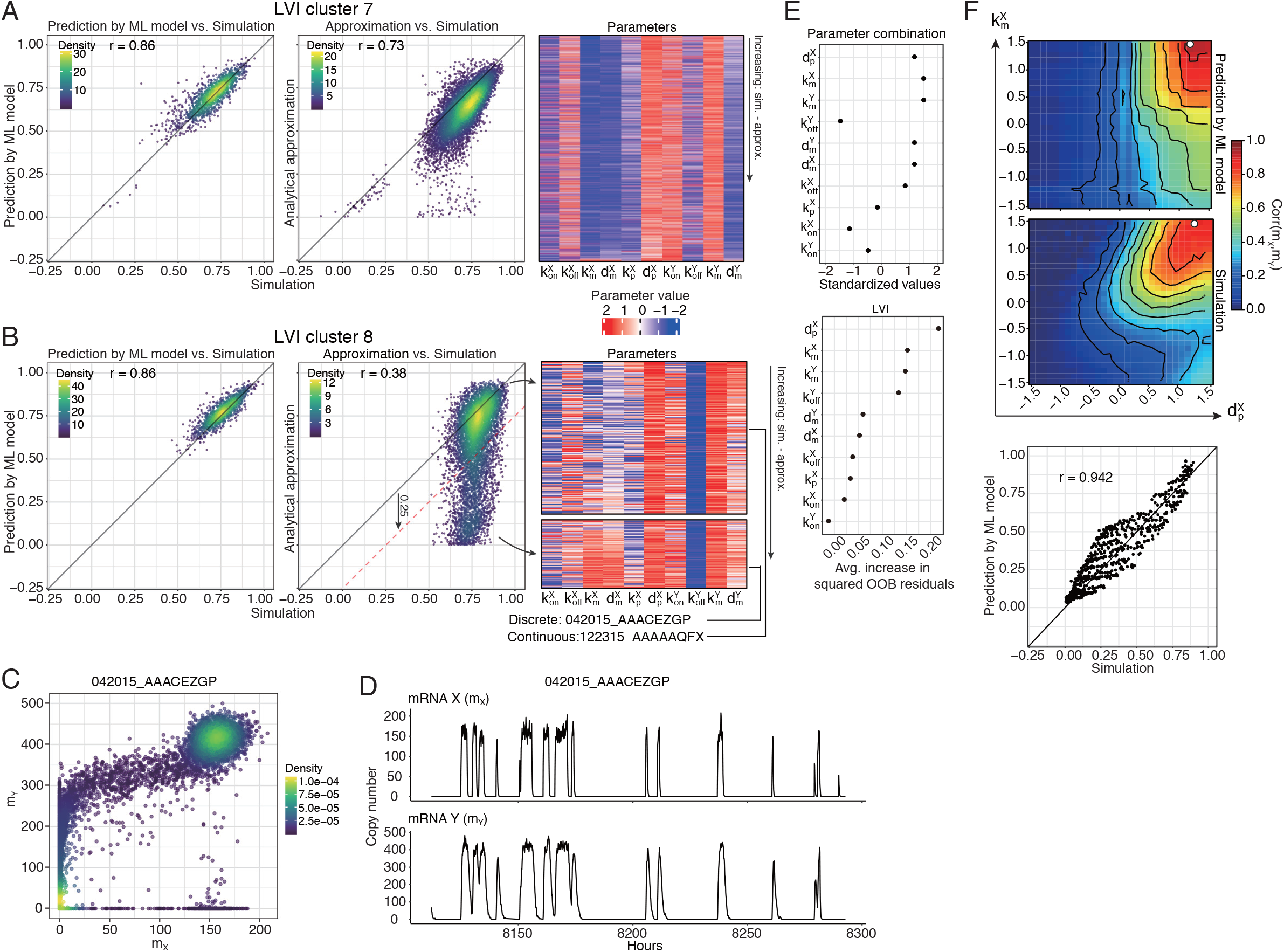
Local variable importance reveals parameter configurations giving rise to coordinated (i.e., both genes), discrete (on or off) stable states that were not predicted by analytical analysis. **(A and B)** Scatter plots showing *Corr*(*m_X_, m_Y_*) of ML model prediction vs. simulation (left) and analytical approximation vs. simulation (center) and heatmap of parameter combinations (rows) (right) in LVI clusters (A) 7 and (B) 8. The parameter combinations in the heatmap were ranked in increasing order by the “error” made by the analytical model compared to actual simulations, i.e., by the difference of *Corr*(*m_X_, m_Y_*) between simulation and the analytical model; for LVI cluster 8 (in 2B), the parameter sets were split into those with differences ≤ 0.25 (top) or >0.25 (bottom). The color scale in the scatter plots denotes the distribution density reflecting the relative abundance of data points (see Methods for details). **(C and D)** (C) Scatter plot and (D) corresponding time trajectories of mRNA X (*m_X_*) and mRNA Y (*m_Y_*) for a representative parameter combination (parameter key: 042015_AAACEZGP) from the bottom heatmap in 2B where the analytical approach performed poorly compared to the ML models. **(E)** The specific parameter combination (the “starting point”) selected for *in-silico* perturbation experiments and its corresponding LVI. Avg. – average; OOB – out-of-bag. **(F)** Contour maps depicting the predicted (top) and simulated (bottom) phenotypic values ((*Corr*(*m_X_,m_Y_*)) as parameters 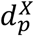 (x axis) and 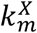 (y axis) were perturbed starting from the selected parameter combination shown in (E) (denoted by white dots); A scatter plot of the predicted vs. the simulated data points from these maps is shown at the bottom.

The LVI can be used to guide fast, high-resolution *in silico* explorations of how parameter perturbations of different extents may affect phenotypes, which can be computationally slow and resource intensive when full-blown simulations are used. For example, the LVI of the parameter configuration discussed above that resulted in discrete correlation (parameter key: 042015_AAACEZGP; Figures 2C and 2D; *Corr*(*m_X_,m_Y_*)=0.83) predicted that the degradation rate of protein X 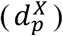 and the transcription rate of gene X (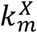 —regulating the “burst” size) were the most important determinants of *Corr*(*m_X_, m_Y_*), while the on-rate of promoters X and Y (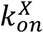 and 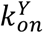) were the least important (Figure 2E). Indeed, as confirmed by actual simulations, tuning 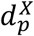 and 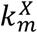 locally affected *Corr*(*m_X_, m_Y_*) substantially, while changing 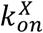 and 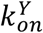 did not (Figures 2F and S3E). Together, our results illustrate that predictive ML models linking parameter and phenotypic spaces can be built successfully and are useful as computationally efficient and phenomenological solutions of the associated SMEs. Our analyses also provided insights on PoV regulation that go beyond those from analytically tractable approaches.

Gene networks containing feedforward interactions are found across phylogeny and biological processes (*20, 21*) (Figure 3A). To assess the ability of MAPPA to analyze more complex networks, we applied it to study two types of feedforward circuits: 1) the coherent type (PPP), in which X positively regulates Y and Z, and Y also positively regulates Z, 2) the incoherent type (PNP), in which X activates both Y and Z, while Y represses Z (*22*). The PPPs can function as delayed activators to filter out transient fluctuations in upstream signals, while the PNPs can serve as accelerated activators or detectors of changes in the input signal over time (*22, 23*). However, the function of these circuits, especially that of the Y arm, is not well understood when stochasticity is present—e.g., how is information transmission from gene X to gene Z (*Corr*(*m_X_, m_Y_*)) regulated by gene Y? We thus defined the phenotype of interest as the ratio (or “fold change” (FC)) between the *Corr*(*m_X_, m_Y_*) of the network with and that without the Y feedforward arm when all the other parameter values remained fixed (Figure 3A; see Methods).

**Figure 3.**
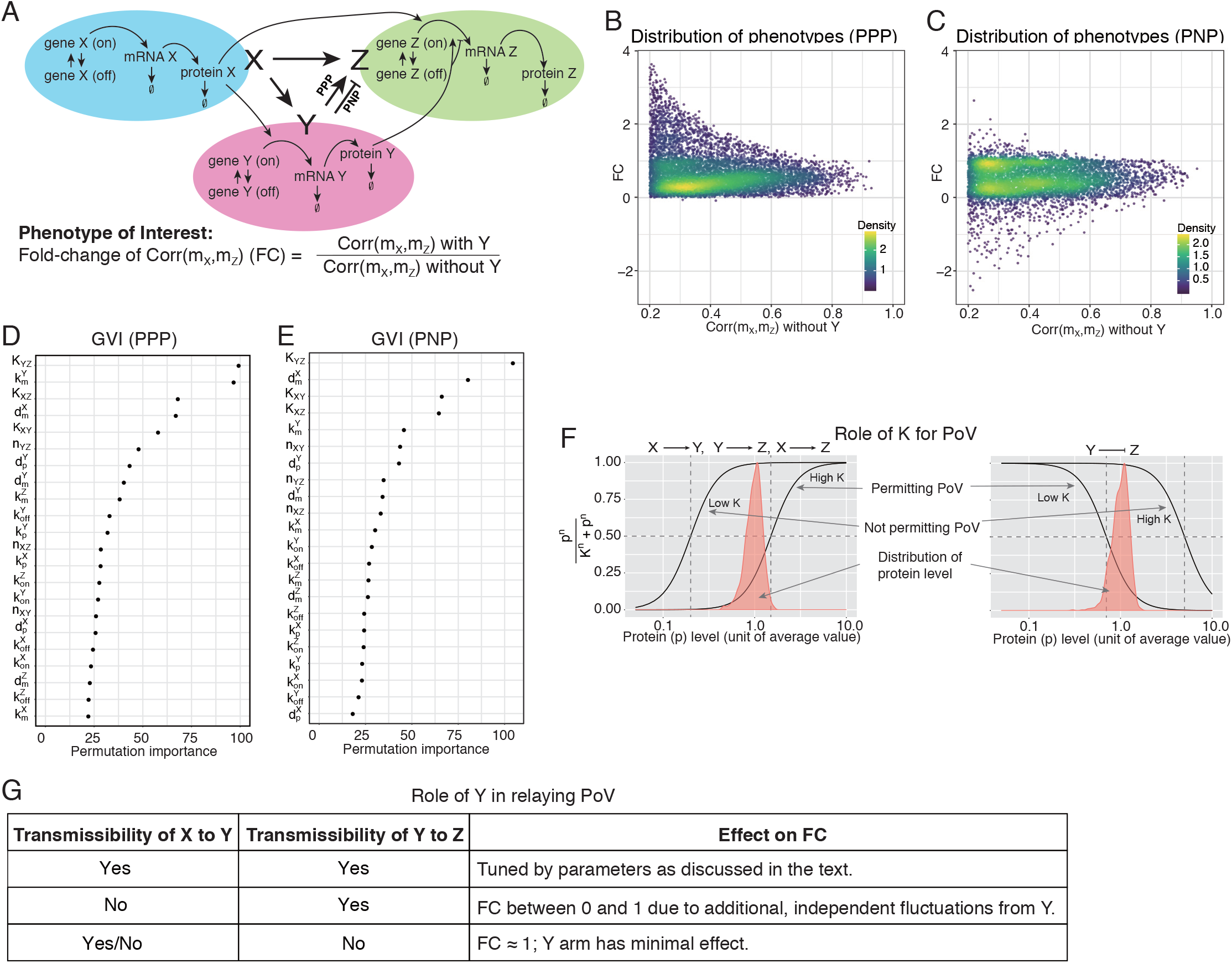
Variability propagation in three-gene feedforward networks. **(A)** Description of the three-gene feedforward network models and definition of the phenotype of interest. The PPP (coherent) and PNP (incoherent) network types are considered. The quantitative phenotype of interest is the ratio of *Corr*(*m_X_, m_Z_*) (or foldchange (FC)) between the network with and without the Y-mediated feedforward; the goal of the analysis is to assess the function/effect of the Y feedforward arm. **(B and C)** Scatter plot of phenotypes, FC versus *Corr*(*m_X_,m_Z_*) without Y for (B) PPP and (C) PNP; each dot corresponds to a single parameter combination sampled. Note that only parameter combinations with *Corr*(*m_X_,m_Z_*) > 0.2 without Y are considered since FC is less robust and would diverge when *Corr*(*m_X_,m_Z_*) without Y is near 0. The color scale denotes the distribution density reflecting the relative abundance of data points (see Methods for details). **(D and E)** GVI of the Random Forest ML model of FC for (D) PPP and (E) PNP. Permutation GVIs are shown. See Figures S5D and S5F for both permutation and impurity GVIs and a comparison between the two. **(F)** Illustrating the role of *K* (*K_XY_*, *K_YZ_*, and *K_XZ_*), the level of upstream input needed to attain half maximal activation of the downstream gene, plays in tuning the propagation of variability/information. The input (upstream protein level) is illustrated as a distribution to depict variability over time within a single cell (or cell-to-cell variation at a given timepoint.) (red), this together with the relative value of K determine whether upstream variations are buffered or transmitted to effect downstream transcription. Left and right panels illustrate a positive and negative regulatory relationship, respectively, between the upstream gene and its downstream target gene. **(G)** Illustrating the main qualitative scenarios of variability propagation in the Y arm and the corresponding effect on the FC phenotype.

Stochastic simulation revealed a notable difference in FC between the two types of feedforward circuits (Figures 3B-C and S4A-B). In PPP, the Y-mediated loop can either increase (FC>1) or decrease (FC<1) *Corr*(*m_X_, m_Y_*), but the correlation was reduced (FC<1) in PNP for most parameter combinations, even down to negative values in some cases (i.e., the direction of correlation flipped); these were more apparent when the correlation was lower (e.g., less than 0.4) when Y is absent (Figures 3B-C). Moreover, PPP and PNP also differ in the time lag needed to maximize the correlation (*Corr*(*m_X_, m_Y_*)): the Y-mediated arm tends to lengthen or shorten the delay between X and Z in the PPP or PNP, respectively (Figures S4C-D), which are consistent with the aforementioned functions of PPP and PNP as delayed and accelerated activators, respectively. Together, simulations revealed distinct functions of the Y arm in regulating information propagation in the PPP and PNP circuits: in a “co-activating” circuit (PPP), the Y arm can increase the correlation extended over longer timescales, while the negative regulating Y arm in the PNP can reduce the transmission of variation from X to Z, thereby maintaining Z homeostasis and reducing Z’s “memory” on X fluctuations.

What regulates FC is less clear. We therefore trained RF ML models that maps parameters to FC (Figure S5A; Methods). These models showed excellent prediction performance (r = 0.93 (PPP) and r = 0.91 (PNP)) (Figures S5B-C). Interestingly, despite their phenotypic differences, the most important parameters for determining FC were similar between the two circuit types (Figures 3D-E, S5D, and S5F). These parameters fall into three categories regulating the transmission of the fluctuations from: 1) X to Y (*K_XY_* and 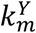), 2) Y to Z (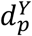 and *K_YZ_*), and 3) X to Z (*K_XZ_*). Most notably, tuning the *K* parameters (Equations 29, 35, and 36 in Methods) can shift the transfer function in and out of the range of variation of the upstream regulator: when operating out of range (at the saturating regime of the transfer function), variation in the activity level of the upstream factor are buffered and thus cannot be transmitted to the downstream gene (Figure 3F). Posttranslational modification of the upstream regulator and epigenetic modification of the promoters/enhancers of the target gene are capable of regulating K in this manner (*24*). Thus, given permitted transmission of variation directly from X to Z, MAPPA pointed to several means for Y to influence the correlation between X and Z (and thus the FC) (Figure 3G).

Similar to the example above (Figure 2), the LVI map revealed that the contribution of individual parameters to FC depends on the parameter configuration (Figures S5E and S5G; see Methods). For example, the Hill coefficients (e.g., *n_YZ_*), which govern the “steepness” of the transfer functions, are generally not important (Figures S5D and S5F). However, LVI showed that *n_YZ_* is important at a particular parameter configuration (parameter key: 011416_AAAAFJWZ) in the PPP circuit (Figure S5H). With this configuration information can be transmitted from X to Z and from Y to Z, but that between X and Y was minimal because *K_XY_* and 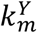 were low and FC was therefore low (0.26) due to the added noise transmitted from Y to Z (Figures S5H-I). Actual simulation confirmed that decreasing *n_YZ_* leads to decreased noise transmission from Y to Z (Figure 3G) and thus an increase in FC (Figure S5J). These data illustrate that by employing even simple feedforward architectures, cells can attain additional flexibility in tuning the co-variation between circuit components (X and Z). For example, having separate “modulatory” Y feedforward arms can be useful when X is a master regulator of many genes: each Y can independently tune information transfer between X and a specific set of downstream genes.

Negative feedback circuit motifs are ubiquitous in biology; their functions include maintaining homeostasis, buffering fluctuations, and driving oscillatory behaviors (*25, 26*). While autoregulatory feedbacks have been analyzed in the context of stochastic gene expression (*27*), the two-gene negative feedback circuit (Figure 4A and Methods), an extension of the simple two gene circuit we analyzed earlier (Figure 1), has received less attention despite its intriguing, implicated roles in generating oscillatory behaviors such as circadian rhythm (*28*). Stochastic simulations of this circuit revealed parameters that resulted in oscillatory behavior (Figures 4A and S6A; Methods). In addition, deterministic modeling and bifurcation analysis confirmed that this circuit is capable of oscillations (limit cycle oscillations or damped oscillations) (Figure S6B and Methods) (*29, 30*). When stochasticity is considered, oscillations (due to “stochastic resonance”) can occur even under parameter regimes that are predicted to not oscillate according to deterministic models (*31–33*), thereby suggesting that in this circuit, stochasticity together with the appropriate coupling (i.e., governing the PoV) between genes X and Y can give rise to oscillations beyond classic “limit cycle” mechanisms.

**Figure 4.**
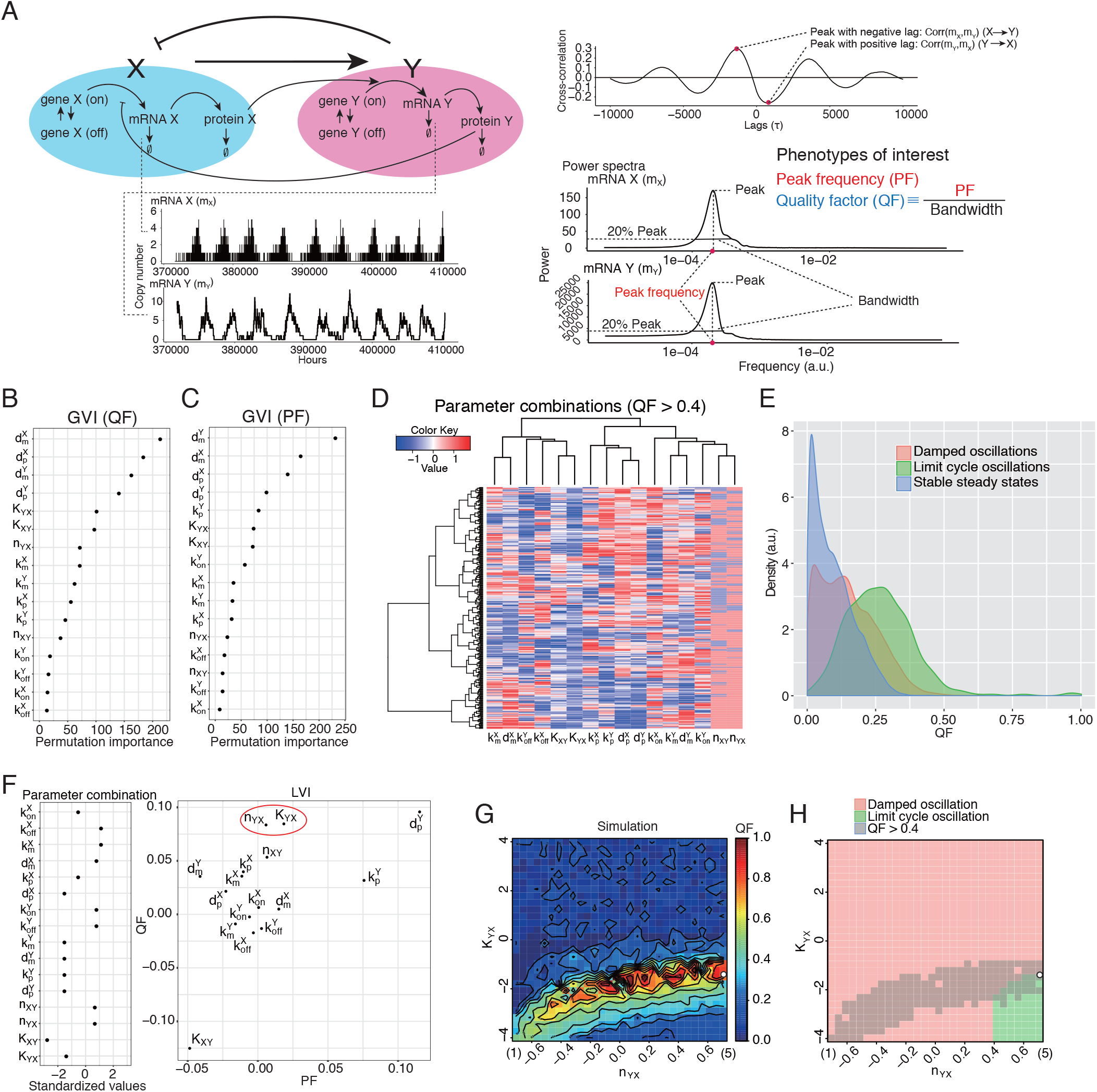
Information transmission and oscillatory behavior of a two-gene negative feedback network. **(A)** The two-gene negative feedback network model and the definition of the phenotypes of interest. Due to the bidirectional regulation between X and Y, peak cross-correlation can be considered with both positive and negative time lags (see Figure S6A). The phenotypes of interest for analyzing oscillatory behaviors include the quality factor (QF) and the peak frequency (PF) of the power spectra in the frequency domain of gene expression dynamics. Example trajectories of mRNAs X and Y are shown for a parameter combination exhibiting oscillations. **(B and C)** GVI of the RF regression model for (B) PF and (C) QF. Permutation GVIs are shown. See Figures S6G and S6H for both permutation and impurity GVIs. **(D)** Hierarchical clustered heatmap of parameter combinations (rows) with QF > 0.4. The color scale denotes the relative (z-score scaled) magnitude of the parameter value. **(E)** Distributions of QF obtained by stochastic simulation for each type of circuit behaviors classified by deterministic ordinary differential equation modeling; DO: damped oscillations, LC: limit cycle oscillations, and SS: stable steady states. **(F)** A selected parameter combination for local, in silico perturbation analysis (left) and its LVI for QF and PF (right; shown as a scatter plot). The two parameters being perturbed are circled in red. **(G)** Stochastic simulation result of QF as *n_YX_* and *K_YX_* are perturbed starting from the parameter combination (denoted by a white dot) shown in (F). Here *n_YX_* is varied continuously between the scaled/standardized values of −0.707 (corresponding to an original Hill coefficient of 1) and 0.707 (original Hill coefficient of 5). The color scale denotes QF. **(H)** The oscillatory behaviors predicted by deterministic modeling in the same parameter space shown in (G) (see Methods). Damped oscillation regions are depicted in pink and limit cycle oscillation regions are in green. The “oscillatory” region (defined as those with QF>0.4) predicted by stochastic modeling is shown in grey.

We thus applied MAPPA to explore how the PoV between X and Y can regulate oscillatory behaviors when stochasticity in gene expression is present. To quantify oscillatory phenotypes, time-varying gene expression levels were transformed to the frequency domain via power spectra analysis. Since a dominant and narrow peak at a specific frequency is expected if the system is oscillating with a relatively constant period and amplitude (Figure 4A, Methods), we focused on two quantitative phenotypes: 1) the quality factor of oscillation (QF), quantifying how tall and narrow the dominant peak is, and 2) the peak frequency (PF), where the peak is located in the power spectrum (Figures 4A and S6C-D; Methods) (*34*).

Predictive ML models for both QF (r = 0.77; Figure S6E) and PF (r = 0.94; Figure S6F) could be built (Methods), although our model tended to underestimate QF when QF is large. Based on GVI, degradation rates (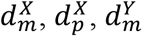, and 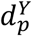) were the most important for predicting both QF and PF (Figures 4B-C and S6G-H), although their relative importance can, again, vary depending on the parameter configuration (Figures S6I-J). Thus, having the appropriate combination of relaxation time scales is required for robust oscillations and for setting the period. Parameter combinations associated with higher QF (QF>0.4) tended to have matching protein degradation rates for X and Y (i.e., 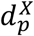 and 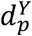 are similar), yet matching mRNA degradation rates (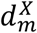 and 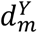) was not necessary (Figure 4D). *K_YX_* also needs to be low so that even low levels of the Y protein can have a sizable suppressive effect on the transcription rate of gene X (Figure 4D). Both *K_XY_* and *K_YX_* (the TF activity needed to achieve half maximal transcription rate of the target gene) and the Hill coefficient (*n_YX_*) of the negative feedback also had high GVI for QF (Figure 4B). Deterministic modeling similarly suggested that a high *n_YX_* is required for oscillations (*35*), suggesting that, in general, parameters like *n_YX_* may regulate both the “average” (as revealed by deterministic models) and fluctuation induced oscillatory phenotypes.

We next asked how stochasticity and PoV shape oscillatory phenotypes beyond the behavior predicted by deterministic models. Deterministic modeling predicted three non-overlapping classes of parameter configurations with distinct behaviors: 1) limit cycle oscillations (LC), 2) damped oscillations (DO), or 3) stable steady states (SS)). We assessed the QF for each of the parameter combinations that fell within these individual phenotypic classes (Figure 4E). This analysis revealed that even the non-oscillatory parameter regimes (based on deterministic modeling) can have non-zero, sizable QF once the effect of stochasticity is considered, and each class has distinct but overlapping distributions of QF. As expected, LC had the largest fraction of its parameter combinations with high QF while SS had the least. More surprisingly, each of the three distributions span wide ranges and even some of the SS parameter combinations can have QF approaching the median of the LC distribution. These results suggest that stochasticity and PoV between X and Y may together be exploited by cells to attain and finetune oscillatory behavior beyond that predicted by deterministic considerations alone (Figure 4E).

To assess the regulatory effects of the key parameters predicted by our ML model, we chose a parameter combination (parameter key: 111315_AAAAENMF) with high QF and varied both *K_YX_* and *n_YX_*, as suggested by their high LVI for QF at this particular point in parameter space (which belongs to the LC class exhibiting relaxation-type oscillations due to the strong feedback (low *K_YX_* and high *n_YX_*) based on deterministic modeling) (Figure 4F) (*35*). Our ML model predicted that increasing *K_YX_* or decreasing *n_YX_* can lower QF, which was confirmed using data from actual stochastic simulation (Figures 4G and S6K). While deterministic modeling suggested a qualitatively similar requirement of low *K_YX_* for limit cycle oscillation (green area in Figure 4H), here with stochasticity considered a higher QF (e.g., QF>0.4) can be attained even when *n_YX_* is lower (grey area in Figure 4H), especially when compensated by a lower *K_YX_* (i.e., higher sensitivity to X suppression) (Figures 4G and 4H). Our analyses thus revealed interesting insights on the regulation of noise induced oscillation in this feedback circuit. By utilizing naturally arising stochasticity and PoV, this circuit could generate oscillation even when *n_YX_* is low and thus far from the DO-LC bifurcation point (where the system transitions from DO to LC; the boundary between green and pink areas in Figure 4H). Operating at a lower *n_YX_* may be biologically more desirable as it can potentially improve operational robustness (e.g., under changing environments) because random switching between DO and LC is less likely (*35*). These results further illustrate how MAPPA can complement and provide information beyond analytically tractable models in a computationally efficient manner.

MAPPA generates and utilizes massive data from mechanistic simulation of biological models and builds ML models to map parameters to phenotypes. The resulting ML models can enable computationally efficient exploration of large parameter spaces and reveal which and how parameters affect a system’s behavior at both the global and local (parameter dependent) levels. MAPPA can guide which parameters to measure, dissect the robustness and “optimality” of the system, suggest evolutionary trajectories, and empower synthetic biology (*3, 36, 37*). MAPPA can in principle be applied to study systems comprising hundreds and thousands of parameters. Given the enormous parameter space, however, it may be computationally intractable to sample sufficient representative parameter combinations for training generalizable ML models. One strategy worth further testing, as we had explored above, would be to start with sparse random sampling and then increase the sampling depth incrementally but biasedly. For example, we can iterate between ML model evaluation (i.e., prediction performance) and targeted sampling from parameter regions associated with desired phenotypes but poor prediction performance. This scheme may converge towards informative models relatively quickly even when the number of parameters is large. For example, Loyola et al. (*38*) suggested an iterative sampling strategy that theoretically does not depend on the dimensionality of the input space, thus effectively avoiding the curse of dimensionality. As future work, incorporating such approaches to MAPPA can potentially enable efficient analysis of networks with orders of magnitude more parameters. Utilizing the large amounts of data generated from bottom-up, mechanistic computational simulation of dynamical systems and the ability of modern machine learning approaches to “compress” such data to generate computationally efficient and interpretable models is a promising direction for dissecting complex dynamical systems.

## Author contributions

KP and JST designed modeling, simulation and data analysis strategies with inputs from TP and YL; KP performed modeling, simulation and data analysis with help from TP and YL; KP and JST wrote the manuscript with inputs from other authors; JST conceived the study and the conceptual framework and supervised the research.

## Acknowledgements

We thank Manikandan Narayanan, Andrew Martins, and Bastian Angermann for valuable inputs and discussions; Ronald Germain and Hao Yuan Kueh for comments on the manuscript; this research was supported by the Intramural Program of the National Institute of Allergy and Infectious Diseases.

---

**Figure S1.**
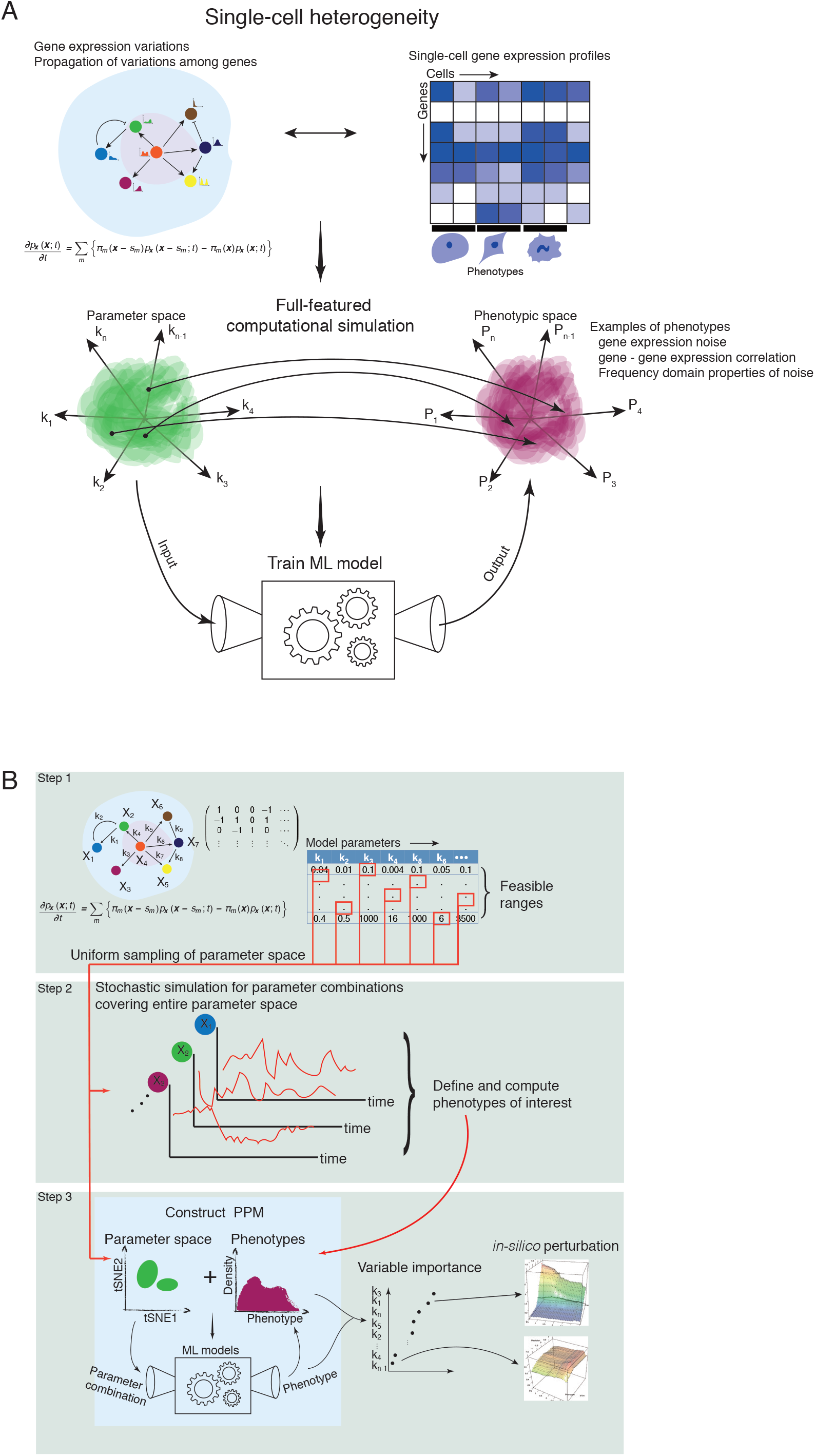
Motivation and conceptual framework. **(A)** Cell-to-cell gene expression heterogeneity is prevalent and can propagate from gene to gene across the gene regulatory network, giving rise to patterns of expression heterogeneity in the cell population that are potentially associated with cellular phenotypes. Given a biological network, our framework samples parameter combinations, conducts stochastic dynamical simulations to generate time series data sets from the network, builds machine learning (ML) models that connect parameter and phenotypic spaces, such as linking parameter values to cell-to-cell gene expression variations and gene-gene correlations across single cells. The ML models can be thought of as phenomenological solutions of the equations governing the stochastic dynamics of the network. They enable much faster computation of quantitative phenotypes from parameter combinations than using full-blown simulations and a better understanding of how the system’s phenotypes are shaped by the parameters. **(B)** The MAPPA framework. Step 1: Model the system as a network of interacting molecular species; their interactions are governed by kinetic parameters; **Step 2**: Uniformly sample parameter combinations from the plausible parameter space and conduct mechanistic/stochastic simulation on each of the combinations and compute the quantitative phenotype(s) of interest from the simulation results; **Step 3**: Construct Parameter-Phenotype Maps (PPMs) by training ML models using the simulated dataset generated in the previous step; PPMs map parameters to phenotypes. The trained ML models can be tested using additional simulated data from parameter combinations distinct from those used to generate the training set. This process can be repeated to improve the PPM, for example, by increasing the representation of parameter combinations that lead to rare phenotypic values. The resultant ML models can be used to explore, in a computationally efficient manner, how parameter perturbations may change the phenotype and delineate which parameters contribute to controlling the phenotype, both globally throughout the parameter space or locally at specific neighborhoods of the parameter space.

**Figure S2.**
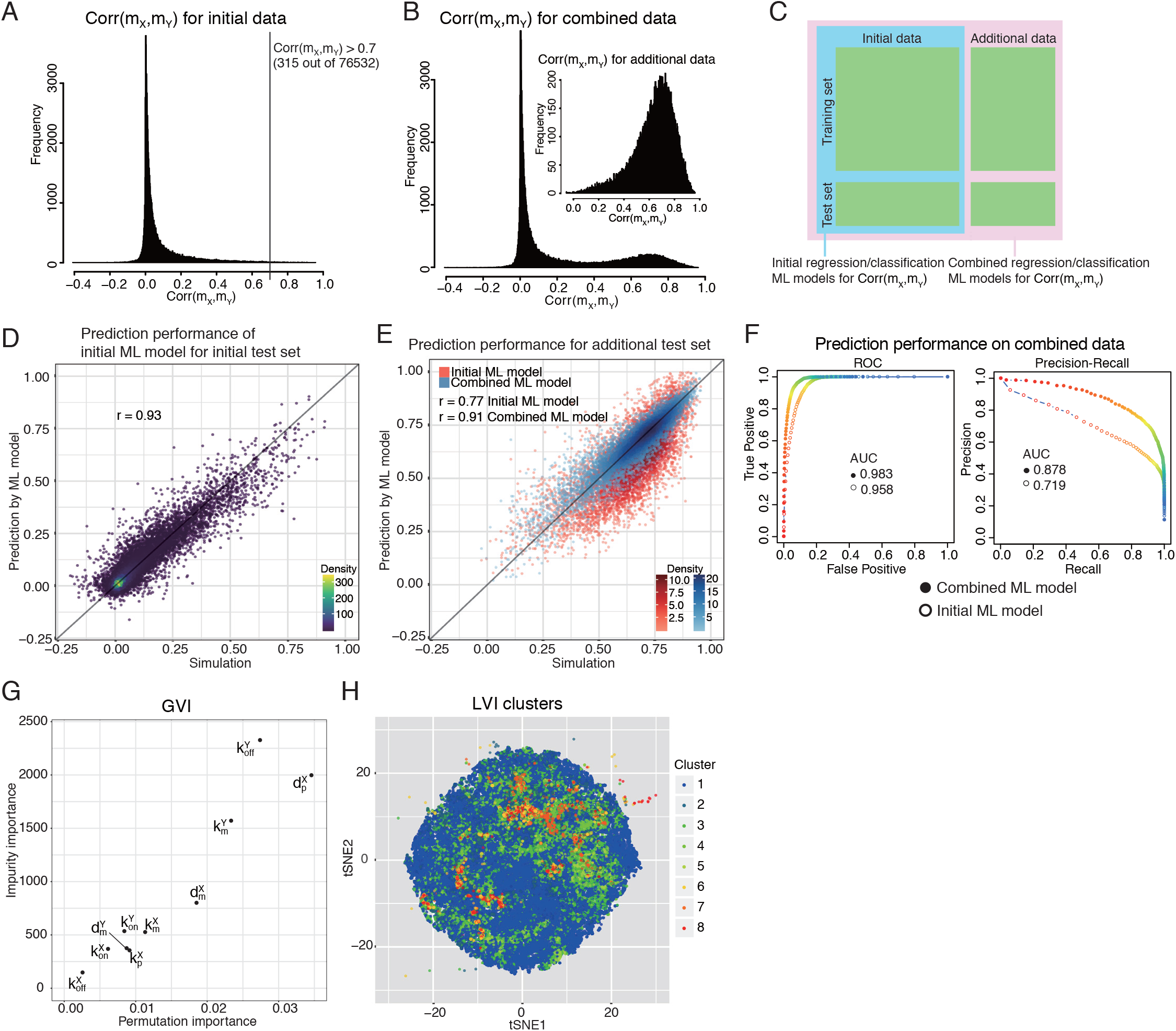
(related to Figure 1) Simulation results and ML models for a two-gene network. **(A)** Distribution of *Corr*(*m_X_, m_Y_*) from the simulations using the first, initial set of sampled parameter combinations (see Figure 1D left panel). Among the 76,532 parameter combinations simulated, only 315 had *Corr*(*m_X_, m_Y_*) > 0.7 with most exhibiting very low correlations. **(B)** Additional simulations were performed on parameter combinations sampled near those shown in (A) with high correlations. Here the distribution of *Corr*(*m_X_, m_Y_*) from both the initial and additional parameter combinations is shown; the inset shows that of the additional parameter combinations only. **(C)** The ML model training-testing scheme involving the initial (within the blue box) and additionally sampled (pink box; outside the blue box) data as described in (B). Each of the two data sets were divided into independent, non-overlapping training and testing sets. We trained 4 ML models (*Corr*(*m_X_, m_Y_*) ~ kinetic parameters (R notation)): two of which were RF regression models, one using the initial (blue box) and the other using the combined (pink box) training set; similarly, two RF classification models for categorical outcomes (high vs. low *Corr*(*m_X_, m_Y_*)) were trained (see Methods). Categorical labels for the classification models: ‘high’ if *Corr*(*m_X_,m_Y_*) > 0.7 or ‘low’ if *Corr*(*m_X_,m_Y_*) ≤0.7. **(D)** Scatter plot showing the predicted (y) vs. the simulated (x) phenotypic values. The RF regression model trained on the initial training set was used to predict the phenotypic value for the initial test set. The color scale denotes the distribution density reflecting the relative abundance of data points (see Methods for details). **(E)** Same as (D) but showing the prediction performance of the RF regression models trained on the initial training set (red) and the combined training set (blue) in predicting the additional test set (enriching for parameter combinations with high *Corr*(*m_X_, m_Y_*)). The color scale denotes the distribution density reflecting the relative abundance of data points (see Methods for details). **(F)** Prediction performance of the RF classification models (*Corr*(*m_X_, m_Y_*) > 0.7 vs. ≤ 0.7) evaluated using the combined test set, as indicated by the receiver operating characteristic (ROC) and precision-recall curves. Hollow circles correspond to data from the RF model trained using the initial training set, while solid circles are data from the RF model trained using the combined training set. (G) Impurity (y axis) vs. permutation (x axis) GVIs for the RF regression model trained on the combined training set are shown together in a scatter plot. **(H)** tSNE plot of all (initial and additional) parameter combinations colored by the cluster ID defined in Figure 1H.

**Figure S3.**
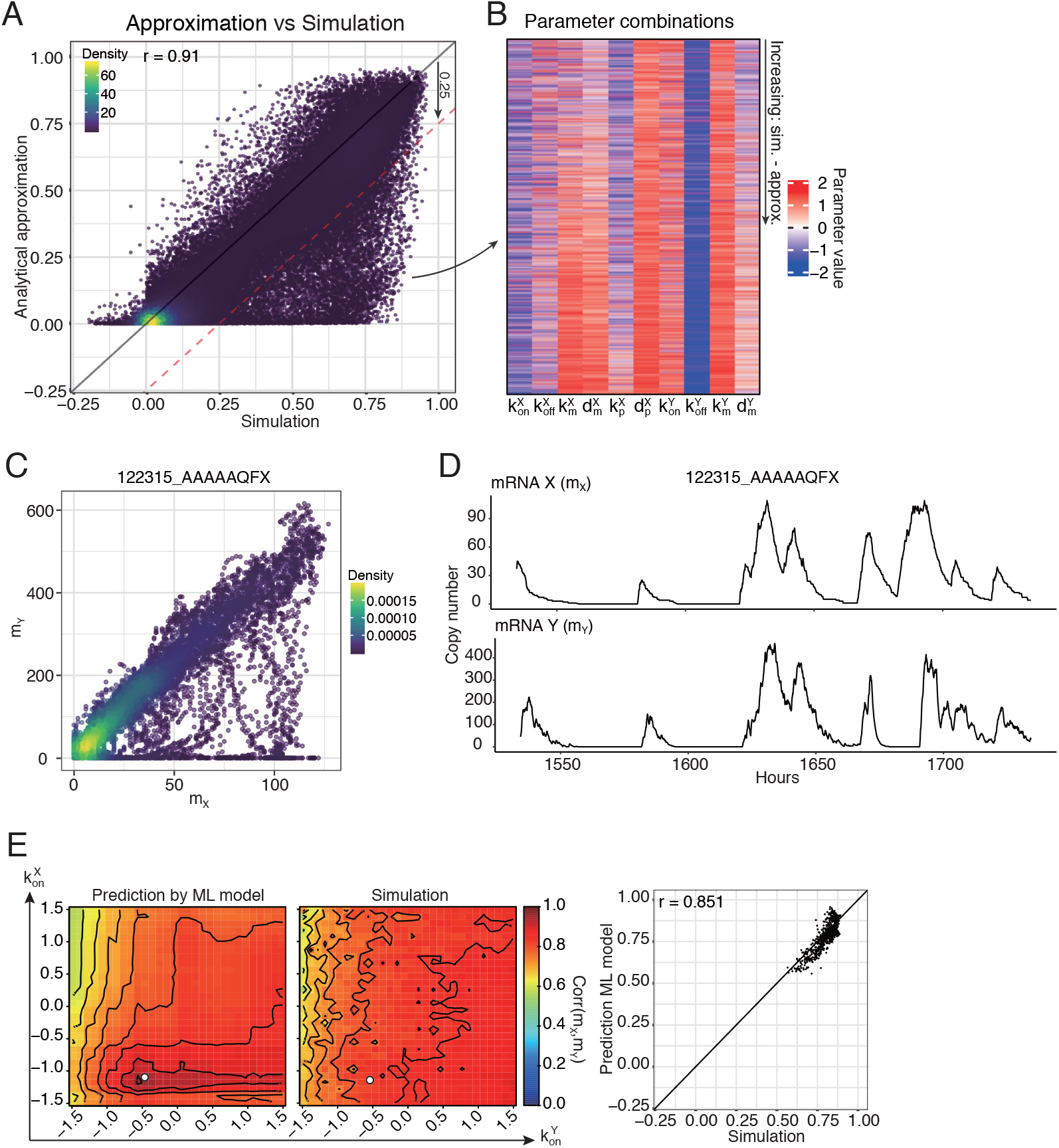
(related to Figure 2) Comparison between ML model predictions and analytical approximations and in silico perturbation analysis of a two-gene network. **(A)** Scatter plot of *Corr*(*m_X_,m_Y_*) computed from analytical approximation (y axis) versus that from stochastic simulation in the entire dataset (both “initial” and “additional” – see Figure S2C). The color scale denotes the distribution density reflecting the relative abundance of data points (see Methods for details). **(B)** Heatmap of parameter combinations (rows) for which the analytical approximation deviate significantly from the simulation results (i.e., with the differences > 0.25 in *Corr*(*m_X_,m_Y_*)) The rows are sorted in increasing order of the difference of *Corr*(*m_X_,m_Y_*) between simulation and analytical approximation. **(C and D)** (C) Scatter plot and (D) corresponding time trajectories of mRNA X (*m_X_*) and mRNA Y (*m_Y_*) for a parameter combination (parameter key: 122315_AAAAAQFX) from the upper heatmap in Figure 2B. **(E)** Similar to Figure 2F but for perturbing 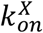 and 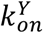 (parameters with lowest LVI) starting from the parameter combination (denoted by white dots) shown in Figure 2E.

**Figure S4.**
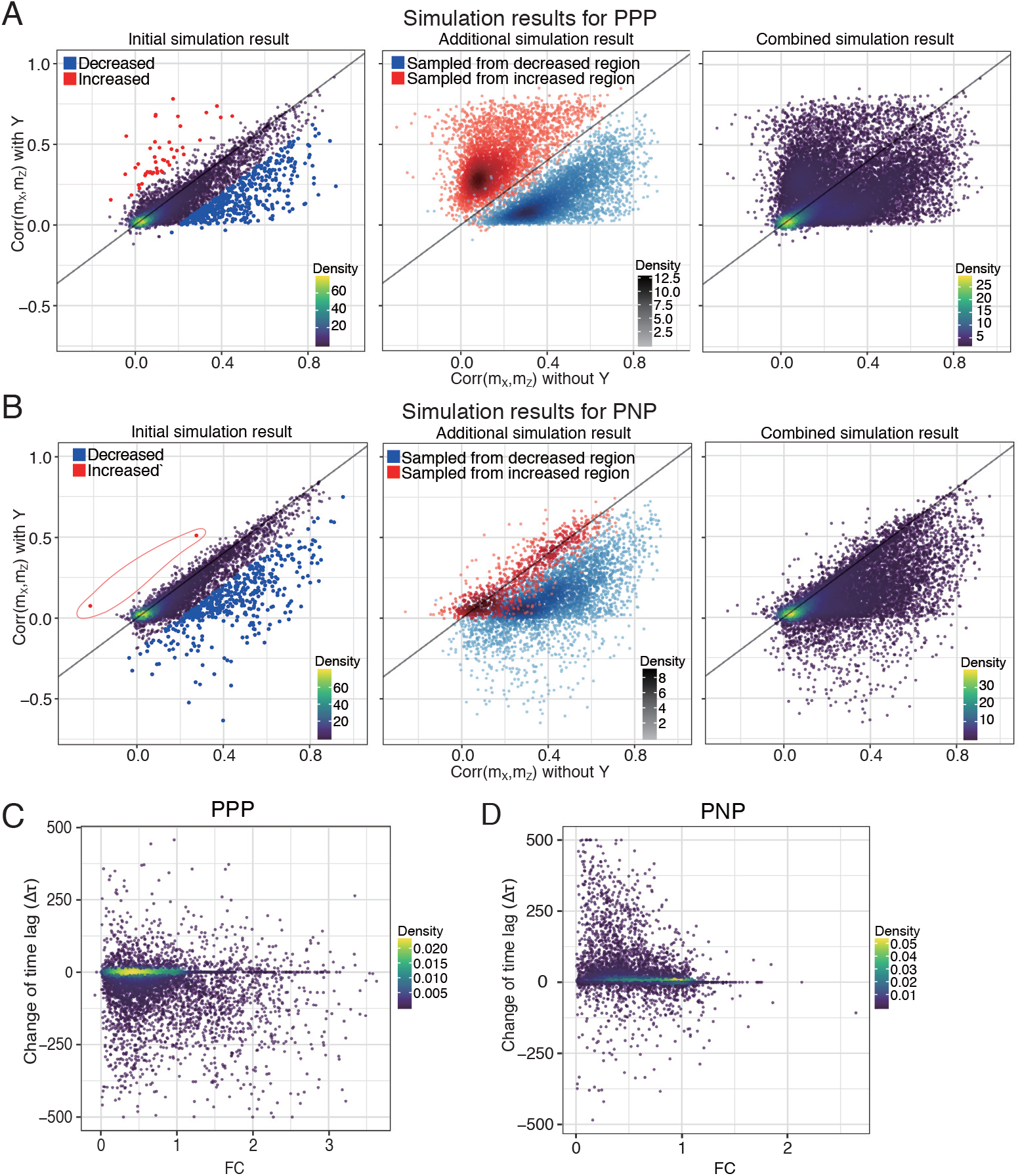
(Related to Figure 3) Simulation results for three-gene feedforward networks. **(A and B)** Simulation results for (A) PPP and (B) PNP circuit types. (Left) Scatter plot showing the simulation results with (y axis) and without (x axis) the feedforward component (mediated by Y) for different parameter combinations (using the initial samples). Those with a difference in *Corr*(*m_X_,m_Y_*) of greater than 0.2 are labeled with red (increased with Y) or blue (decreased with Y). (Center) Simulation results for the additional parameter combinations. The red (blue) dots correspond to parameter combinations sampled around those that resulted in the red (blue) dots in the left panel. (Right) The same but showing both the initial and additional parameter combinations. The color scale denotes the distribution density reflecting the relative abundance of data points (see Methods for details). **(C and D)** Differences in the lag time (y) for achieving peak cross-correlations between the circuits with and without Y for (C) PPP and (D) PNP. PPP tends to lengthen time lags (Δ*τ*, < 0) while PNP tends to shorten time lags (Δ*τ*, > 0). For PNP, only parameter combinations with positive *Corr*(*m_X_,m_Y_*) with Y were included for proper comparison between PPP and PNP. The color scale denotes the distribution density reflecting the relative abundance of data points (see Methods for details).

**Figure S5.**
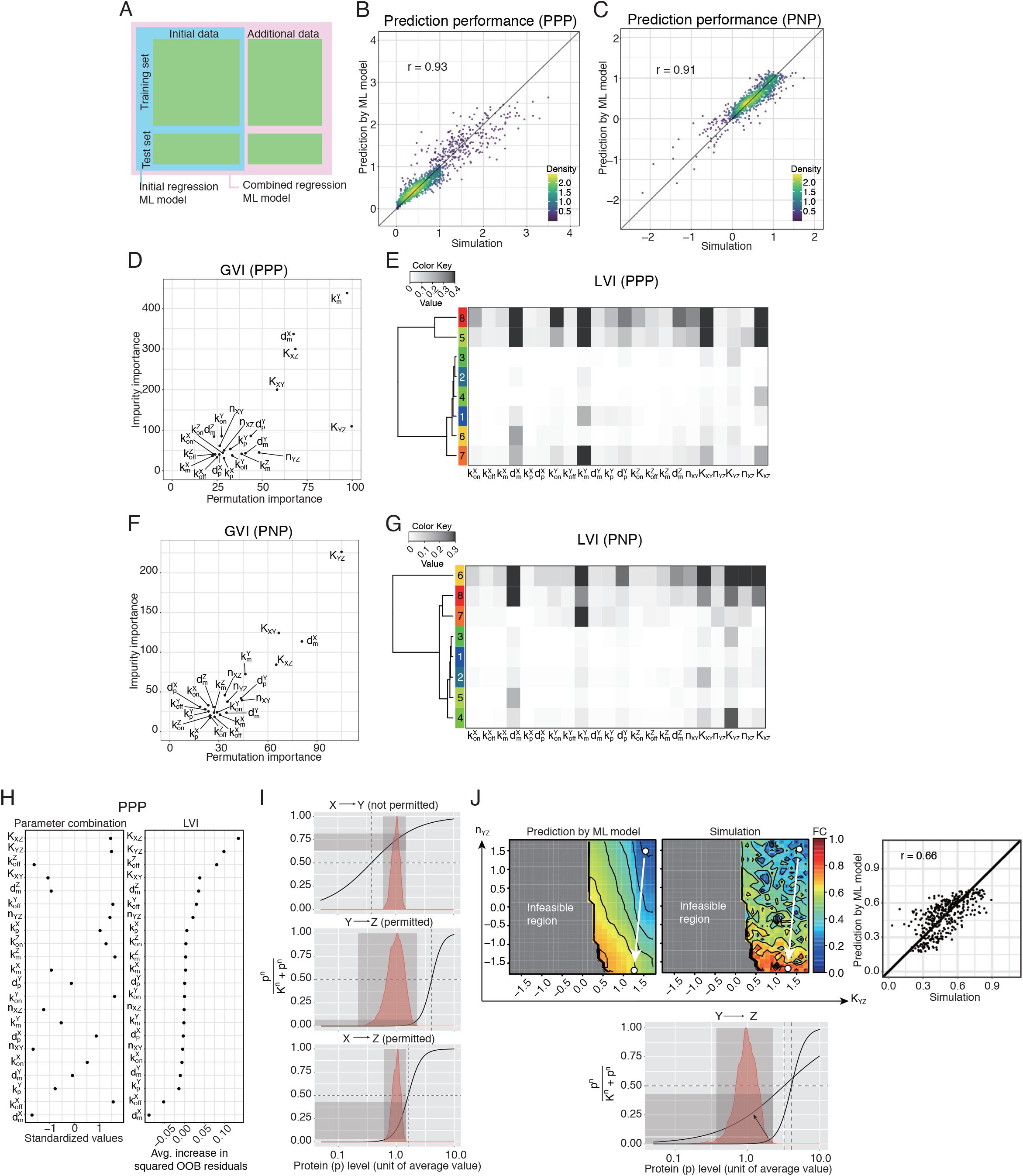
(Related to Figure 3) ML models for three-gene feedforward networks. **(A)** ML model training scheme (similar to Figure S2C). We trained RF regression models on the initial and the combined (initial and additional) training set for both network types and assessed prediction performance using the initial and the combined test set, respectively. For the rest of the analyses, including obtaining GVI and LVI and in silico perturbation experiments, ML models were trained using the combined training set. In the main figures we reported results from the combined dataset. **(B and C)** The prediction performance of the RF regression model for FC trained using the combined training sets for (B) PPP (*r* = 0.93) and (C) PNP (*r* = 0.91). The color scale denotes the distribution density reflecting the relative abundance of data points (see Methods for details). **(D and F)** Impurity and permutation GVIs are shown together for (D) PPP and (F) PNP. Ranks for highly important parameters are largely consistent between the impurity and permutation importance measures. **(E and G)** The LVI (from the ML model for FC) of the parameter combinations were clustered and the average values of each cluster is shown for (E) PPP and (G) PNP. The cluster number is shown in the color bar (Methods). **(H)** The specific parameter combination from PPP selected for local perturbation analysis, and its LVI (right panel). Avg. – average; OOB – out-of-bag. **(I)** Distributions of protein levels (*p_x_, p_y_*, and *p_z_*) (red) and the transfer functions (black curve – see Figure 3F) between the indicated upstream protein and downstream gene at the selected parameter combination. The shaded areas indicate the effective regulation regimes between the input (upstream protein) and output (transcription rate). **(J)** Prediction (by RF model) and validation (based on stochastic simulations) for the perturbation on the two indicated parameters (x and y axes) starting from the selected parameter combination shown in (H). (Top) Prediction and simulation of phenotypic values (the two contour maps on the left), and a scatter plot comparing prediction and simulation for the given perturbations (right panel). The grey regions in the contour maps represent parameter combinations deemed biologically infeasible (Methods). The starting point (white dot) of the arrow is the parameter combination shown in (H), and the arrow indicates the shift in the transfer function (especially in the Hill coefficient) arriving at the second parameter combination (the end point of the arrow) as a result of the parameter perturbation; the bottom plot (similar to (I)) shows the changes in the transfer function between these two parameter combinations (indicated by the arrow) and how the second parameter combination allows higher transmission of variation (higher FC).

**Figure S6.**
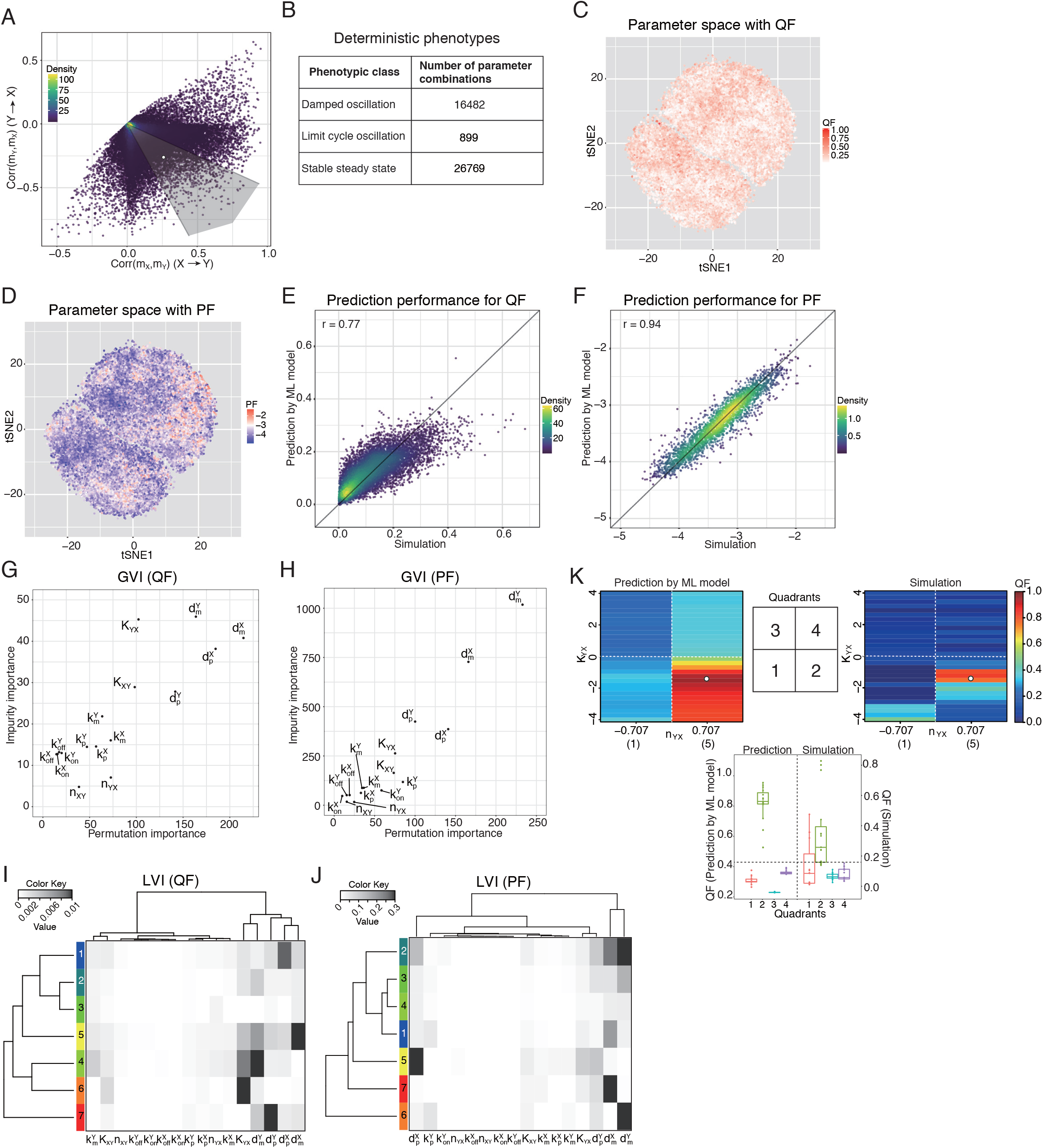
(related to Figure 4). Simulation results, nonlinear bifurcation behaviors, and ML models of the two-gene negative feedback network. **(A)** Simulation results showing peak cross-correlations between *m_X_* and *m_Y_* with negative time lag (x axis – *Corr*(*m_X_, m_Y_*); information flows from X to Y) vs. positive time lag (y axis – *Corr*(*m_X_, m_Y_*); information flows from Y to X) for the sampled parameter combinations. Shown in the shaded region are those parameter combinations with comparable magnitudes of *Corr*(*m_X_, m_Y_*) and *Corr*(*m_Y_, m_X_*) (defined as within twofolds between the positive- and negative-lagged correlations), which potentially exhibiting oscillations. The example oscillatory trajectories shown in Figure 4A was generated from the parameter combination denoted by the white dot. The color scale denotes the distribution density reflecting the relative abundance of data points (see Methods for details). **(B)** Number of parameter combinations for each type of oscillatory behavior predicted by deterministic modeling; damped oscillations (DO), limit cycle oscillations (LC), and stable steady states (SS). **(C)** tSNE plot of the sampled parameter combinations colored by QF. **(D)** tSNE plot of the sampled parameter combinations colored by PF. See Methods for additional details on the logged unit of PF. **(E)** Scatter plot of predicted vs. simulated QF to illustrate the prediction performance of the RF regression model for QF. The color scale denotes the distribution density reflecting the relative abundance of data points (see Methods for details). **(F)** Scatter plot of predicted vs. simulated PF to illustrate the prediction performance of the RF regression model for PF. The color scale denotes the distribution density reflecting the relative abundance of data points (see Methods for details). **(G and H)** Impurity and permutation GVIs are compared and shown together in scatter plots for (G) PF and (H) QF. Ranks for the most important parameters are consistent between the two measures. **(I and J)** The LVI (from the RF regression model) of the parameter combinations were clustered and the average values of each cluster is shown for (I) QF and (J) PF (see Methods). The cluster number is shown in the color bar. **(K)** Comparison between ML prediction and stochastic simulation of QF as *n_YX_* and *K_YX_* were perturbed starting from the selected parameter combination (denoted by white dots) shown in Figure 4F. We compared the discrete/qualitative behavior by dividing the space into four quadrants with high/low values for *n_YX_* and *K_YX_* (shown in the middle) since the ML model was trained based on only two possible values of *n_YX_*, −0.707 (on the relative scale; the original Hill coefficient value is 1) and 0.707 (the original Hill coefficient value is 5) (see Methods). The QF predicted by the ML model is shown on the left of the quadrant map and the actual simulation is shown on the right. The change in QF across the quadrants is qualitatively consistent between the simulation and the prediction (bottom plot). The full simulation result, as the two parameters were perturbed along the continuous range, is shown in Figure 4G.

## Supplementary Materials

## Glossary

MAPPA: **MAchine learning of Parameter-Phenotype Analysis** – the name of our framework.
Parameter space: Multidimensional space in which each dimension is defined by the biologically plausible range of each parameter.
Phenotypes: The quantities reflecting specific aspects of dynamical or stationary behaviors of the model/system, which are defined and can be computed using dynamical/stochastic simulation results for each parameter combination.
Phenotypic space: Multidimensional space defined by plausible ranges of each phenotype.
PPM – Parameter-Phenotype Map: Quantitative relationship between parameter values and phenotypes of interest. In our work PPMs are fitted as ML (Machine Learning) models.
GVI – Global Variable Importance: The relative contribution of a parameter for predicting phenotypes averaged over parameter combinations in the training set.
LVI – Local Variable Importance: The relative contributions of a parameter for predicting phenotypes at a particular point in parameter space (i.e., at a specific parameter combination).
SME – Stochastic or Chemical Master Equation: Equations describing the dynamic evolution of probability distribution of system states in chemical reaction networks.
SSA – Stochastic Simulation Algorithm: A simulation algorithm that generates dynamic trajectories of system states (e.g. the abundance of chemical species) in a chemical reaction network. The timing and type of reactions are determined probabilistically based on the current system state and kinetic parameters. Ensemble of time trajectories constitute time evolution of probabilities of states described by SMEs.
PoV (Propagation of variation), information transfer, information propagation, transmission of information, transmission of variation: These terms were used interchangeably; they refer to the phenomenon that variation in gene expression across single cells (or dynamic fluctuations over time within single cells) can be propagated from gene to gene in gene regulatory networks. This can be captured by gene-gene correlations across single cells in a cell population (e.g., via singe cell transcriptomic data).
PPP (see Figure 3A): A *coherent* feedforward circuit motif. X positively regulates Y (P). Y positively regulates Z (P). X positively regulates Z (P).
PNP (see Figure 3A): An *incoherent* feedforward circuit motif– “incoherent” because Z is regulated by X and Y in opposite directions. X positively regulates Y (P). Y negatively regulates Z (N). X positively regulates Z (P).
FC: Fold-change of *Corr*(*m_X_, M_Y_*) between PPP or PNP circuits with and without the Y feedforward arm.
Transfer function: a mathematical (e.g., Hill) function describing how the activity of upstream transcription factors affects that of the downstream promotor/enhancer.
QF – Quality Factor: A quantity measuring the extent of oscillation in a system.
PF – Peak Frequency: The dominant frequency of oscillation in a system identified as the most dominant peak in the power spectra (frequency domain analysis) obtained from dynamic trajectories.
LC – Limit Cycle Oscillations: Persistent oscillations with a well-defined frequency/period.
DO – Damped Oscillations: Transient oscillations followed by settling down to stable fixed states.
SS – Stable Steady States: Evolution to fixed stable states without any transient oscillations.

## Supplementary Text

### Learning the Parameter-phenotype map (PPM) of Stochastic Network Dynamics

Given a stochastic gene network model and associated kinetic parameters, we seek to understand how the phenotypes of the system behave and change across the parameter space. Here, phenotypes can be any quantitative measures assessing certain aspects of the dynamical behavior of the given model. (Figure S1A). While the quantitative relationship between parameters and phenotypes in a given system can be mathematically complex, we hypothesize that the mapping can be captured quantitatively by interpretable ML models such as Random Forests (RF) in which the contribution of individual parameters to achieve accurate mapping can be delineated (*39*). For example, the gene expression dynamics of a network of genes and proteins in cells can be modeled by SMEs (Figure S1A). However, analytical solutions of non-linear SMEs (e.g., those with on-off promoter dynamics) are generally intractable and evaluating how parameters affect a phenotype of interest, e.g., the correlation between two genes over time (Figure S1A), rely on conducting computational simulations over a large number of parameter combinations. Here we sought to use ML models to learn the PPM.

MAPPA comprises three steps (Figure S1B). In step 1, we model a system as a network of interacting entities (e.g., proteins, mRNAs, cells), whose states/levels are governed by stochastic birth-death processes (e.g., transcript production and degradation). While still poorly measured, particularly *in vivo*, plausible ranges of some parameters can be obtained from the literature or approximated based on physical and/or biological constraints. Next, in step 2, using methods designed for uniform sparse sampling of highdimensional parameter spaces (Methods), we obtain parameter combinations from the plausible parameter space and conduct stochastic, dynamical simulations of the system using each one of the parameter configuration samples. We then compute quantitative phenotypes of interest (here we focus on correlation of expression between genes in single cells, but the MAPPA approach is applicable to any phenotypes/modeling combinations) to obtain an *in-silico* dataset that links parameter values to phenotypes for the sparse sample of values. In the final and critical step (step 3), we train a ML model that quantitatively maps parameters to phenotypes, and we evaluate the predictive performance of the model by using parameter combinations not used in the training set. Using this approach, we can utilize arbitrarily large training and testing sets with size limited only by computational capacity. Any good ML approach generating interpretable models can in principle be used, here we use RF because it: 1) has good ML performance (*40*), 2) can capture non-linear relationships between parameters and phenotypes, and 3) can delineate parameter importance both locally or globally (i.e., which parameters contribute to phenotypic variations at or independent of a specific location in parameter space, respectively).

Once we have a predictive ML model, we can use it as a “phenomenological” solution of the SME to efficiently predict phenotypes from parameters without using computationally intensive simulations (Figure S1A). Guided by dimensionally-reduced visualizations and information on which parameters contributed to prediction, we can further evaluate the system and test our understanding by *in-silico* perturbation analysis, e.g., by assessing how well we can predict phenotypic changes as the parameter values are altered (Figure S1B). These interactive, exploratory assessments are efficiently enabled by the ML model without full stochastic simulations; they can further help reveal the design principles of the systems and suggest parameter optimization strategies to attain specific phenotypes in synthetic gene circuits (*41*). To illustrate these use cases, we have developed an interactive website to allow the exploration of PPMs we analyzed (https://phasespace-explorer.niaid.nih.gov).

## Methods

### Model Description

Here we describe variables, parameters, and reactions constituting our models in Figures 1A, 3A, and 4A.

#### Model variables

The chemical species (or variables) used in this study are as follows (Figures 1A, 3A, and 4A):

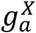: Gene X with an “active” promoter (on-state)
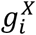: Gene X with an “inactive” promoter (off-state)
*m_X_*: mRNA transcribed from gene X
*p_X_*: Protein translated from mRNA X
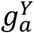: Gene Y with an “active” promoter (on-state)
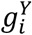: Gene Y with an “inactive” promoter (off-state)
*m_Y_*: mRNA transcribed from gene Y
*p_Y_*: Protein translated from mRNA Y
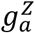: Gene Z with an “active” promoter (on-state)
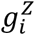: Gene Z with an “inactive” promoter (off-state)
*m_z_*: mRNA transcribed from gene Z
*p_z_*: Protein translated from mRNA Z

#### Kinetic parameters

The range of the kinetic parameters (Table S1) was obtained from the experimental literature (*18, 42–47*), similar to what we did in our previous work (*12*). We summarized the prior experimental findings into biologically feasible ranges for each parameter (Table S1). Based on the reported range of copy numbers of transcription factors (TFs) and correlations between protein copy numbers and other quantities such as translation rate constants (*47, 48*), we obtained modified ranges of the transcription and translation rate constants, which were applied to upstream genes acting as TFs in the models (Table S1).

Over the course of the model development across different network motifs, we tested different ways to specify K (which can be interpreted as the level of the upstream TF needed to achieve the half maximum rate of transcription) when sampling parameter combinations. For the two-gene network (Figure 1), K was fixed to a single value as in our previous work (Table S2)(*12*), while for other circuit motifs (Figures 3 and 4), K was not fixed. For the two-gene negative feedback network (Figure 3), the unit of K was the copy number, which is the same as that for proteins (Table S4). For the three-gene feedforward networks, we tested specifying K as a relative quantity, namely as a ratio to the mean copy number of the upstream TF estimated from the corresponding deterministic model at steady state (Table S3). For downstream analyses including ML model training and visualizations, we decided to make the unit of K consistent across models as the relative ratio, and thus we converted the unit of K in the two-gene negative feedback network copy number to the relative ratio. Observing that the resultant values of K span several orders of magnitude (10^−4^~10^4^), we applied the base-10 logarithm transformation, resulting in K spanning the range −4 to 4.

#### Chemical reactions and the deterministic dynamics for each system

The following biochemical reactions are modeled in the two-gene network (Figure 1B).

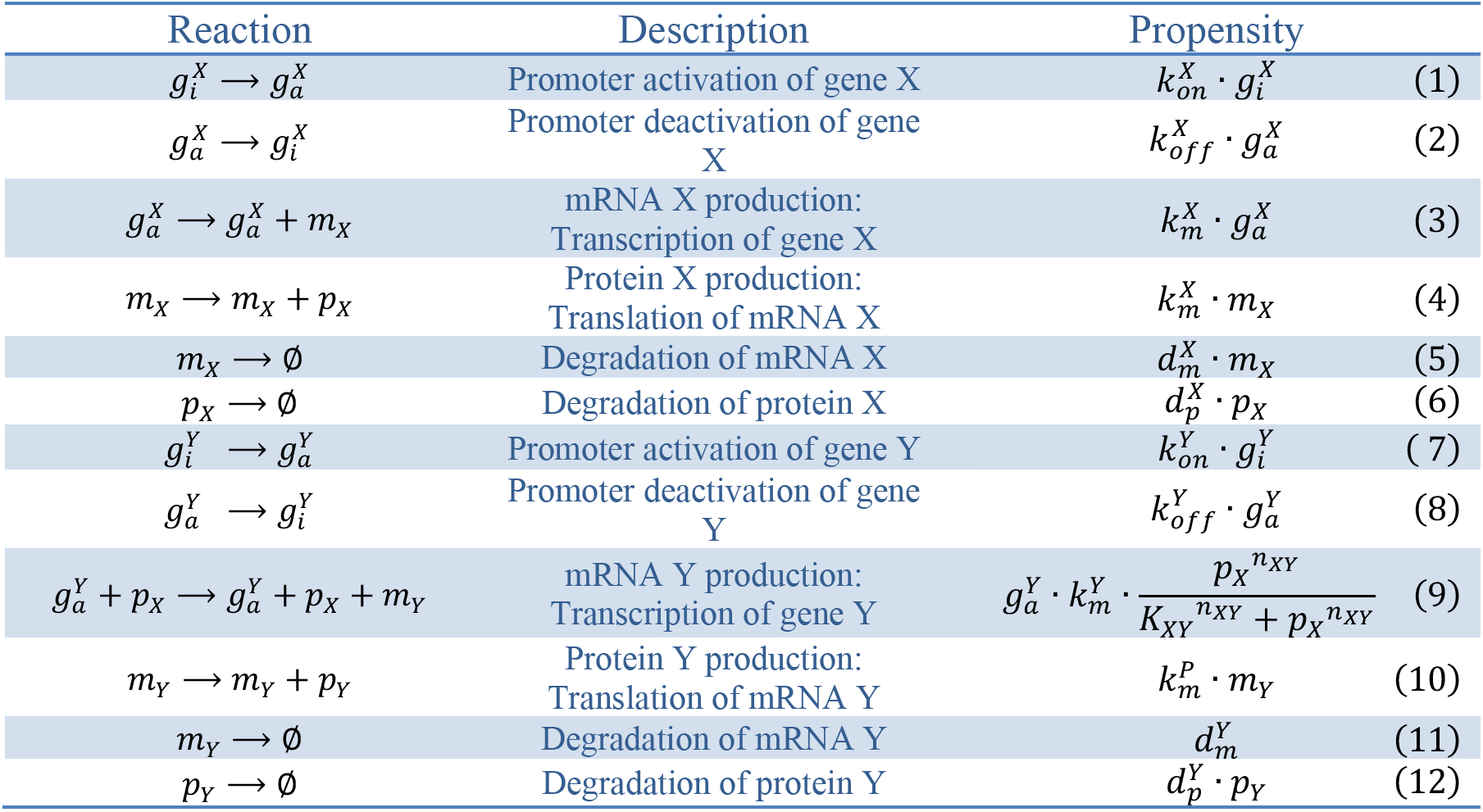

The following equations are deterministic descriptions of the reactions above using ordinary differential equations:

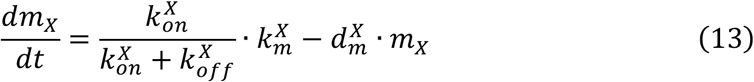

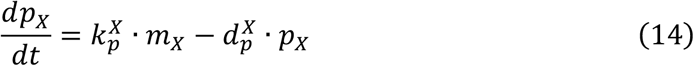

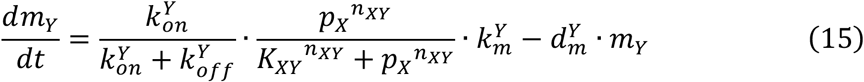

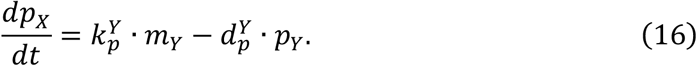

Stationary states can be estimated by setting derivatives in the left-hand side of equations above equal to zero, resulting in the following expressions:

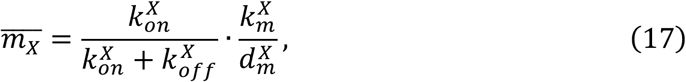

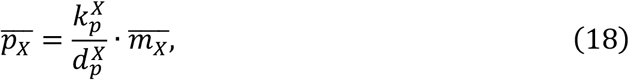

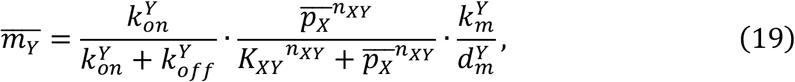

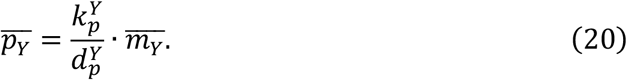

The following reactions describe the elements of the three-gene feedforward networks (PPP and PNP) (Figure 3A).

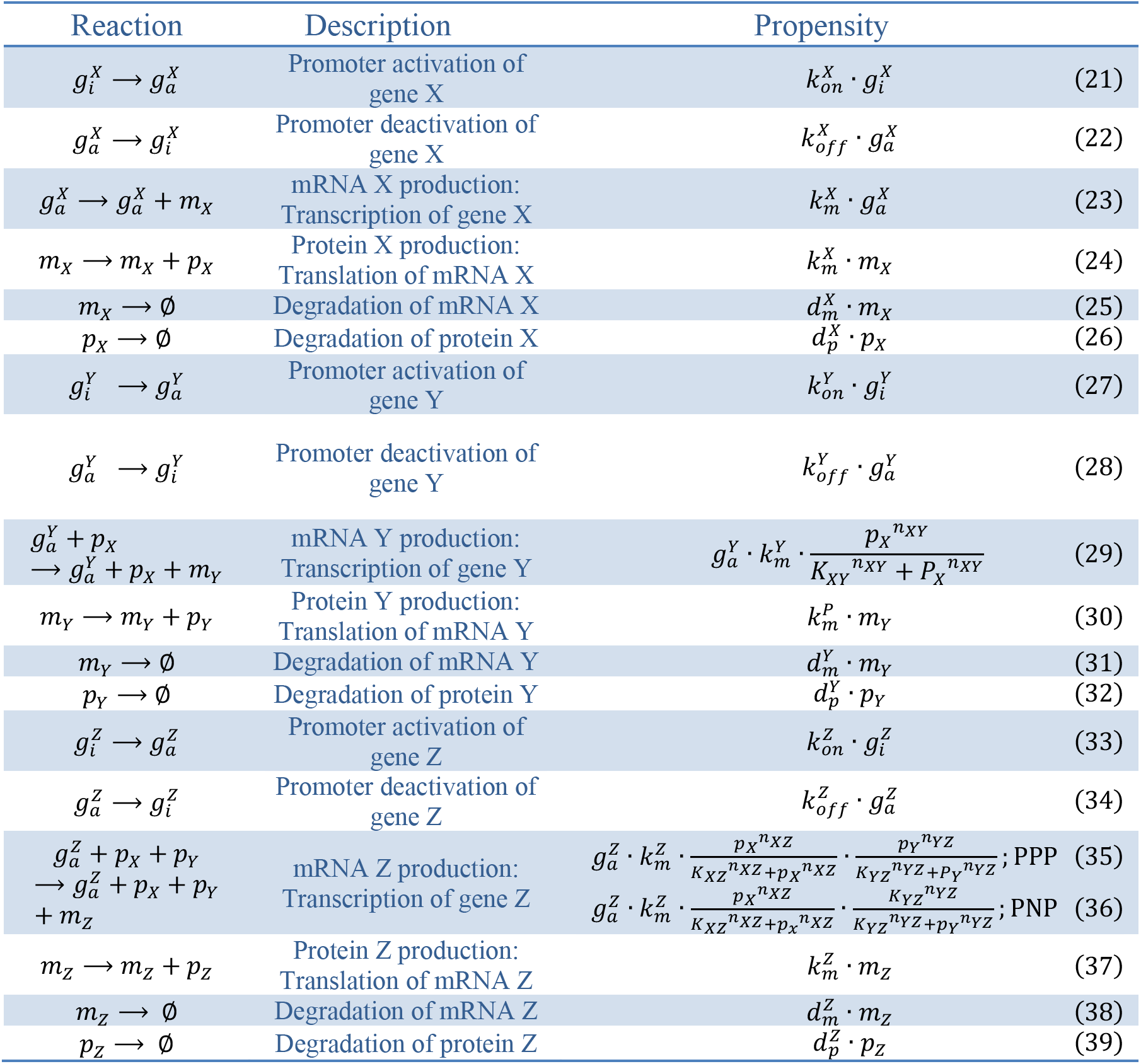

The deterministic descriptions are:

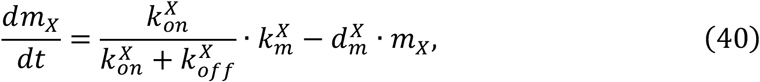

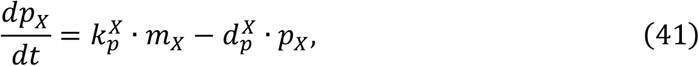

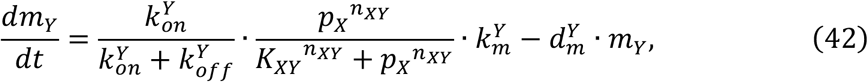

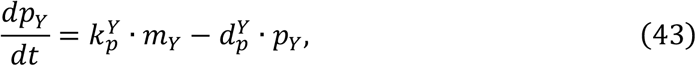

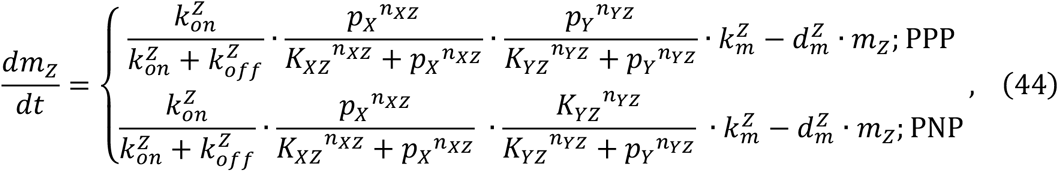

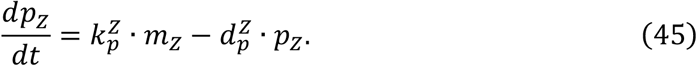

Stationary states were obtained as follows:

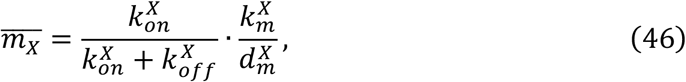

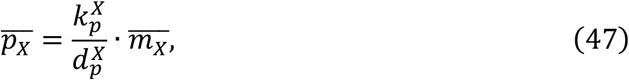

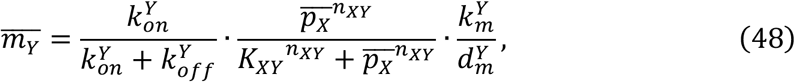

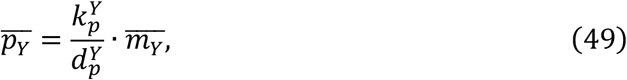

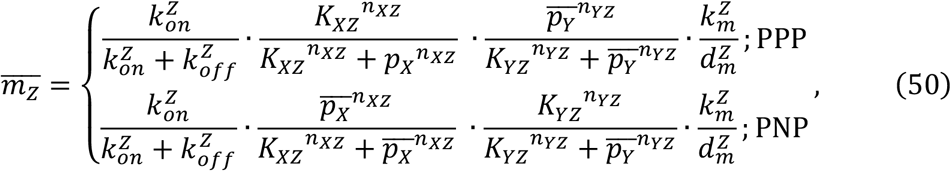

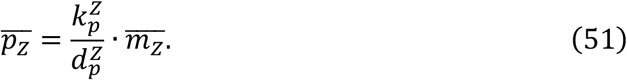

The following reactions describe the elements of the two-gene negative feedback network (Figure 4A).

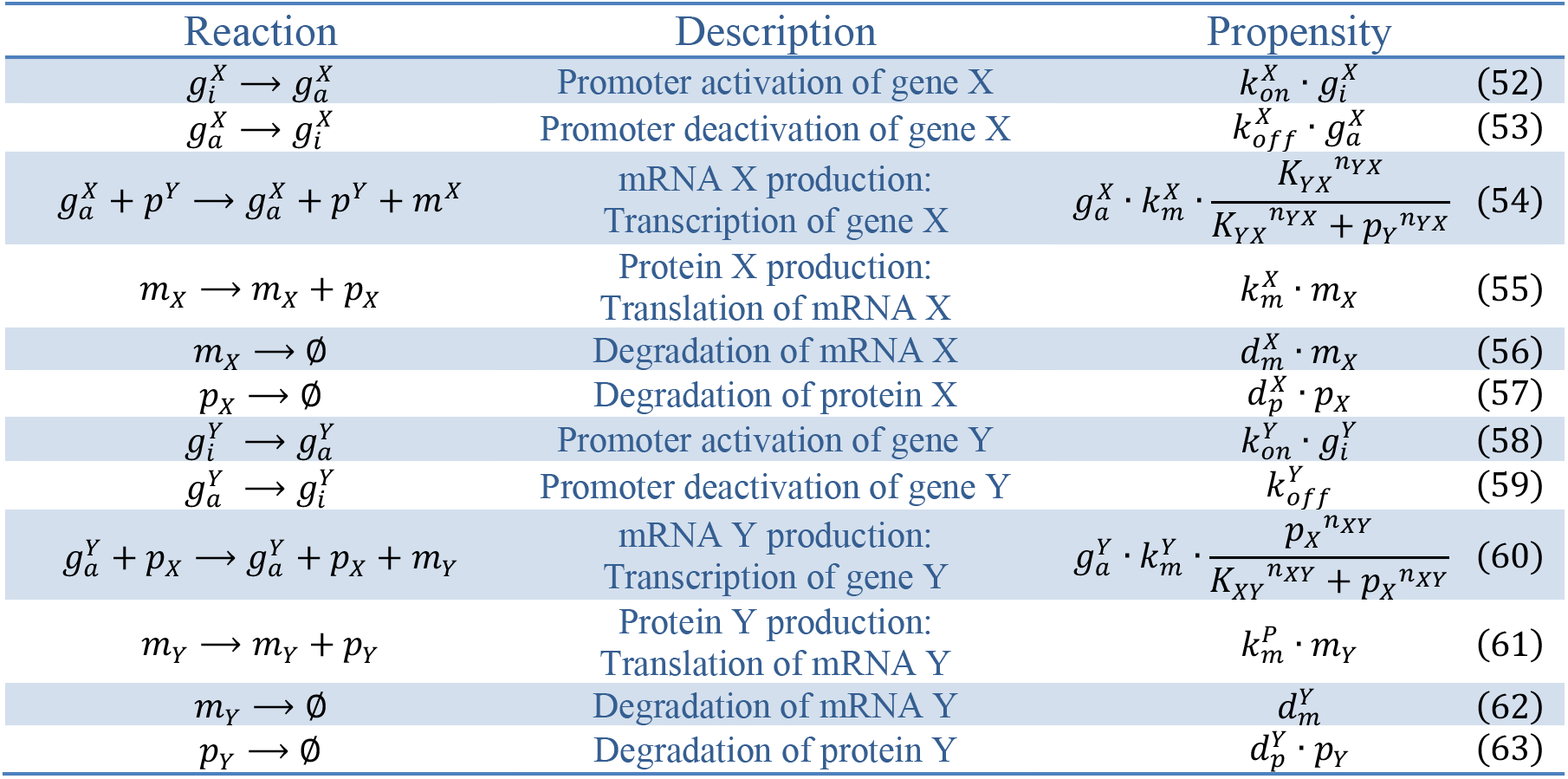

Corresponding deterministic descriptions of reactions are:

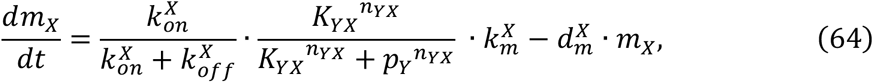

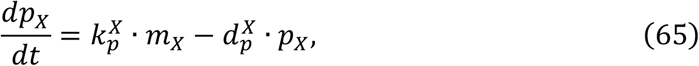

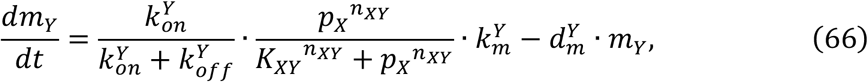

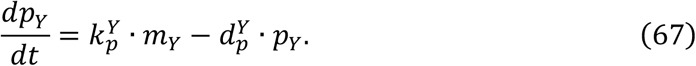

The corresponding stationary states are:

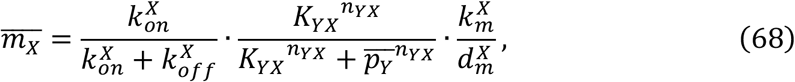

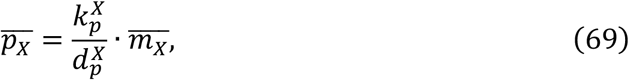

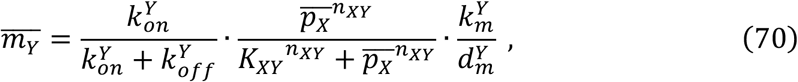

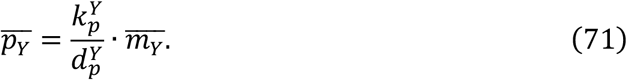

### Sampling of Parameter Combinations

To obtain the system’s phenotypic behavior throughout the biologically plausible parameter space, our strategy was to sample parameter combinations unbiasedly (but sparsely to keep computational cost reasonable).

We first used a simple ‘uniform grid scheme’ for the two-gene network and the two-gene negative feedback network. The range of each parameter was divided into discrete points uniformly in the original or logarithmic scales based on the range of each parameter determined by the distribution reported in the literature (*43, 47*). Then, we sampled parameter combinations by randomly selecting a value out of the grid points for each parameter. For the two-gene network, for example, it has 10^10^ possible parameter combinations since each of the 10 parameters was divided into 10 grid points; there a total of 10^5^ parameter combinations were sampled (Tables S2). For the two-gene negative feedback network having 12 parameters total, 8, 2, and 2 were divided into 10, 5, and 2 grid points, respectively, resulting in a total of 10^14^ possible parameter combinations (Tables S4).

Second, we used the ‘Sobol’ sampling scheme for the three-gene feedforward networks due to the larger number of parameters. The Sobol’ sequence is a low-discrepancy sequence, which fills the unit interval (0,1) more evenly than a pseudorandom sequence (*49*), thus is appropriate for our goal of sampling parameter combinations as uniformly as possible in the high-dimensional parameter space. Using a Sobol’ sequence generator implemented in the *randtoolbox* package in R, a 16-dimensional Sobol’ sequence of 10^5^ points filling a (0,1)^16^ hypercube was generated. Then, this sequence was rescaled linearly in the original or logarithmic scales to fit the parameter ranges previously specified for the three-gene feedforward networks (Table S3). To further specify the network type (out of the 8 possible types (*22*)) for each of parameter combinations, we added a sign (positive and negative) to the Hill coefficients and randomly assigned either a + or – to *n_XY_*, *n_YZ_* and *n_XZ_*, respectively, where the +/− signs represent positive and negative regulation/influence on downstream promoters, respectively. Thus, either activating or repressive Hill functions were used based on the sign of the Hill coefficients during simulations. In the main text/analysis, we only considered two of the most prevalent types (PPP and PNP; Figure 3A).

We also note that Latin hypercube sampling is another possible scheme, although we did not use in our current work (*50*). In the context of global sensitivity analysis, it has been reported that both the Sobol’ sequence and the Latin hypercube work better than pseudorandom sequence, but the relative performance between Sobol and Lain hypercube is less clear (*50–52*).

#### Parameter key

Each of the sampled parameter combination was assigned a unique key to make it easily identifiable. The basic syntax for parameter keys is [date of sampling]_[letter combinations] (e.g., 111315_AAAAENMF).

### Additional Sampling Around Specific Parameter Neighborhoods

Once simulations for the uniformly sampled parameter combinations were conducted for the two-gene and three-gene feedforward networks (Figures 1 and 3) and phenotypes of interest were computed, we sought to obtain a more detailed picture around the regions of the parameter space that exhibit interesting but rare phenotypic states (e.g., high correlations between the genes in the two-gene network). We thus sampled additional parameter combinations around those regions, conducted simulations for these combinations, and augmented this additional data for training new ML models (see Figures S2C and S5A).

The detailed method we implemented is as follows. First, parameter combinations exhibiting phenotypic states of interest were selected. Second, for each parameter combination of interest, the sub-range of detailed sampling for each parameter was created (with 1/10 the width of the original parameter range and centered at its value in the parameter combination). Third, additional parameter combinations were sampled on these sub-ranges using the uniform grid sampling scheme or the Sobol’ sampling scheme as described above. Finally, we conducted stochastic simulations for these new parameter combinations and computed the phenotypes of interest based on the simulation results. Note that for the three-gene feedforward circuit motifs, the parameter combinations for additional sampling were chosen based on a hard cutoff: we selected those whose *Corr*(*m_X_, m_Y_*) in the circuit with Y differ from that without Y by least 0.2 (Figures S4A-B). This was chosen qualitatively through visual inspection of the left panels of Figures S4A and S4B to enrich for parameter combinations in the off-diagonal regions.

### Stochastic Simulation Scheme

As mentioned in the main text, we used Gillespie’s Stochastic Simulation Algorithm (SSA) to generate dynamic trajectories for each of the parameter combinations (*16*). To save space for data storage, we stored the simulation data once every 5 minutes. To ensure that the copy numbers of mRNAs and proteins stay within biologically feasible ranges, we applied a filtering step before starting simulations, such that only parameter combinations for which the estimated mean (steady-state) copy numbers of mRNAs and proteins of genes do not exceed 1,000 and 2,000,000 copies, respectively, can proceed to simulation (*47, 48*).

We used the following procedure to obtain stationary time trajectories. First, we estimated the time scale under which the system would fluctuate around the mean copy number of chemical species based on deterministic differential equations. Each chemical species fluctuates differently in accordance with the “firing” rates of the chemical species and the slowest reactions determine the overall timescale of fluctuation. For proteins and mRNAs, the contributors to the firing rate are degradation and synthesis rates. The mean firing rate can be estimated as:

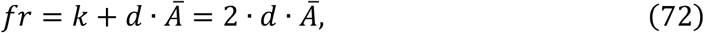

where *fr*: the firing rate, *k*: the synthesis rate, *d*: the degradation rate, *Ā*: the mean copy number of the species; since *k* = *d* · *Ā* in the stationary state, we have an expression that depends only on the degradation rate and the mean copy number. Assuming this firing occurs as a Poisson process, the variance of the number of firing events per unit time is also the firing rate itself. We can define a time scale, *τ_f_* as:

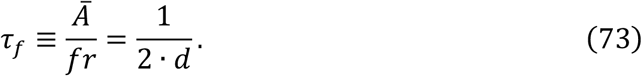

This as a mean time interval during which the variance of the number of firing events is the same as the mean copy number of the chemical species, which can be interpreted as the time it takes to randomize the system such that the “information/memory” about the copy number is lost after this time interval. A related interpretation is that the waiting time of the Poisson process is exponentially distributed with mean waiting time for a single firing event being the inverse of the firing rate (1/(2 · *d* · *Ā*)), thus it would take on average *τ_f_* units of time for *Ā* firing events to occur. For promoters, for example, *τ_f_* can be estimated as 1/*k_off_* or 1/*k_on_*. The maximum of *τ_f_* among all chemical species, *τ_f,max_*, can be considered as a good approximation of the time scale of fluctuation for the system. To obtain a stationary time trajectory of the system, a simulation needs to span multiple intervals of *τ_f,max_* to allow the system to explore distinct regions of the state space. This argument is supported by theoretical studies, for example, see (*14, 53*).

Having defined the time scale of fluctuation, *τ_f,max_*, given system with a particular parameter combination, we divided the simulation into two phases: 1) a burn-in phase with a duration of 4 · *τ_f,max_* and 2) the main phase with duration of multiples of 20 · *τ_f,max_* (see below). The simulation begins with the burn-in phase with the initial condition of zero copies for mRNAs and proteins and inactivated states for promoters, which allows the copy numbers of each species to build up to or near the stationary values; the data from the burnin phase is discarded in later analyses. Then, the main phase begins and generates time trajectories of each molecular species until the stationary test is passed (see below) or a time limit, *t_max_*, is reached. Only the trajectories that passed the stationary test were used for downstream analyses.

The stationary test was conducted after every 20 · *τ_f,max_* and the test was performed using only the last 2/3 of the data/trajectories (partly to mitigate the risk that the copy number did not yet reach near the steady-state values) and consisted of two steps. In the first step, the mean values of two halves of the data (latter 2/3) were compared. If the difference between the two values is less than a cut-off (expressed as a percentage of the mean value of the first half of the data), then the second step is applied. If not, the simulation would continue for another 20 · *τ_f,max_*. In the second step, we used the KPSS test (*54*) to evaluate stationarity for higher order moments since the phenotypes of interest involved cross-correlations and power spectra involving second moments. We stopped the simulation if the generated trajectory passed the test or continued for another 20 · *τ_f,max_* if it failed the test.

### Machine Learning Scheme

We used Random Forests (RF) (*39*) to train machine learning (ML) models that learn the nonlinear relationships between parameters and phenotypes. We combined each parameter combination with its phenotype computed from the simulations (e.g., correlations between mRNAs X and Y) to form a data table; we then partitioned the data into training and test sets with the ratio of ~4:1 and only the training set was used in fitting the model. The function *randomForest* in the R package *randomForest* was applied to the training set to construct an ensemble of 500 decision trees (*39*). Once trained, we tested the ML model using the unseen test set. The prediction performance of the ML model was shown as Receiver Operator Characteristic (ROC) and recall-precision curves for classification ML models (the area under these curves (AUC) was used as the quantitative metric), and the Pearson correlation coefficient between the predicted and simulated values was used for the RF regression based ML models (for predicting continuous values such as gene-gene correlation). For GVI, LVI, and *in-silico* parameter perturbation analyses, we trained ML models by using the entire data set (i.e., combining the training and test sets) for maximal performance since our goal is to predict the effect of unseen perturbations followed by evaluation by additional stimulations.

For the two-gene (Figure 1) and three-gene feedforward (Figure 3) networks, to evaluate whether the use of additional samples drawn nearby the parameter combinations exhibiting desirable phenotypes (see above) can lead to improved prediction accuracy, we first trained ML models using only the “initial” data (without additional samples) and used the “combined” data (with additional samples) to train another model (Figures S2C and S5A). We saw that the performance of the combined ML models was indeed better (or at least comparable to) than that of the initial ML models (Figures S2E-F, data not shown for the three-gene feedforward networks). We thus used combined ML models for the rest of analyses, including the determination of GVIs and LVIs and in the *in-silico* perturbation experiments.

We used RF regression models for continuous value phenotypes. To address the inherent bias of RF regression (*55, 56*), we applied a bias correction method for all ML regression models we trained in this study (*56*). Briefly, we trained additional models for the “error” or the residuals *y – ŷ*, where y is the phenotypic value from the training set and *ŷ* is the value predicted by the uncorrected, original RF regression model. The bias-corrected predictions were obtained by adding the predicted corrections from the residual ML model.

### Variable Importance

The *randomForest* package in R generates variable importance both at the global level (global variable importance (GVI), which can be thought as average over all data points in training sets) and in the local level (local variable importance (LVI)) for each individual data point in the training sets with the ‘importance’ and ‘localImp’ options turned on, respectively (*39, 57*). Two types of global variable importance were computed: permutation and impurity GVIs. The permutation GVI is generated based on the increase in the out-of-bag prediction errors (using the training set only) after randomly permuting each input variable. The impurity GVI is generated by measuring the total decrease in the node “impurity” (which quantifies the heterogeneity/entropy of the outcomes underneath each tree node; Gini index is used for categorical outcome variables and the residual sum of squares for continuous outcome variables) conferred by each variable in the training set. LVI is the increase in the out-of-bag prediction error on a specific data point of the training set (thus a particular point in the parameter space) after permuting each of the input variables. The permutation GVI was generated by averaging the LVIs from all data points. Although GVI gives a general overview, the variable importance can differ across parameter space as captured by the LVI. We grouped the individual LVIs (at each point in the parameter space) by hierarchical clustering and visualized the resulting clusters using tSNE plots, and used them to guide *in-silico* perturbation experiments.

For the hierarchical clustering heatmaps (Figures 1H, S5E, S5G, S6I, and S6J), we showed the average values of LVIs for each cluster rather than showing all LVIs for individual parameter combinations. The number of clusters were chosen based on visual inspection of the original heatmaps to allow visualization of major patterns capturing the full qualitative diversity of the LVIs. Detailed heatmaps showing all LVIs can be generated on our website (https://phasespace-explorer.niaid.nih.gov).

### tSNE Visualization and Embedding of Additional Samples

We used t-distributed stochastic neighbor embedding (tSNE), a technique for dimensionality reduction, to visualize high-dimensional parameter spaces. We first generated a reference tSNE plot in 2D space using the original sampled parameter combinations. Since the tSNE algorithm does not provide a general parameterized transformation from higher to lower dimensional spaces, adding additionally sampled parameter combinations (see above) to an existing visualization requires a new tSNE plot. Thus, we implemented a customized algorithm for embedding additional points in an existing tSNE plot by following Appendix D of Berman et al. *(58)* and applied it for generating Figure 1E. The procedure is briefly summarized below.

Starting with a lower dimensional tSNE embedding, *Y*, of the original high dimensional data, *X*, where *X* and *Y* are matrices, and the rows in them, *x_i_* and *y_i_*, correspond to individual data points with *i* = 1,2,…,*N*, with *N* being the number of data points. Then, for an additional high-dimensional point (with the same dimension as *X*), *z*, we want to obtain lower dimensional embedding, *w* (with the same dimension as *Y*), on the reference embedding, *Y*. To accomplish this, as in the tSNE algorithm, we first defined transition probabilities as

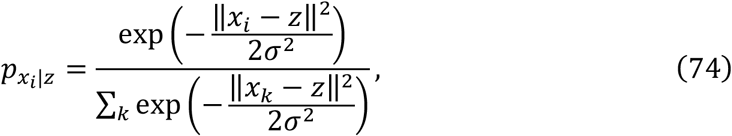

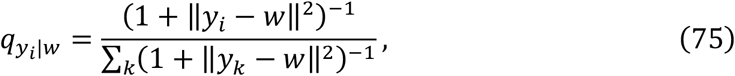

where ǁ⋯ǁ denotes the Euclidian norm of a vector inside, and σ is related to the perplexity, P, a parameter used in the tSNE algorithm, roughly equivalent to the number of nearest points that z can perceive as specified by the following relation,

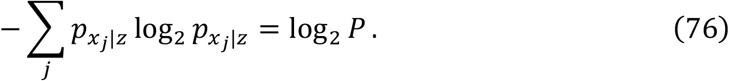

These transition probabilities reflect the similarity between the existing data points and the additional point we would like to embed. Next, the lower dimensional coordinates, *w*, was obtained by minimizing the Kullback-Leibler divergence between, *p_x_i_|z_* and *q_y_i_|z_*, *KL*(*p||q*) by tuning each component of *w* as:

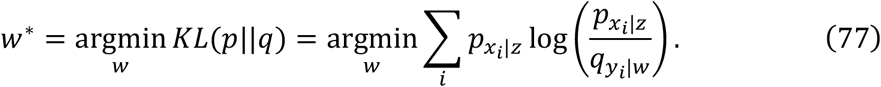

### Analytical linear noise approximation of the correlation between mRNAs X and Y in the two-gene network

We derived an analytical approximation of *Corr*(*m_X_, m_Y_*) by following the approach of Elf and Ehrenberg (*14*) and Paulsson (*17*). With stationary assumption and linearization of the SMEs describing the system, the following relationship (a Lyapunov matrix equation) can be derived,

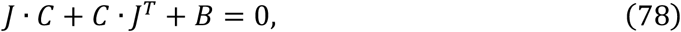

where *J*: the Jacobian of the deterministic equations, *C*: the covariance matrix, and *B*: the diffusion matrix. *J* and *B* at the stationary state were obtained as:

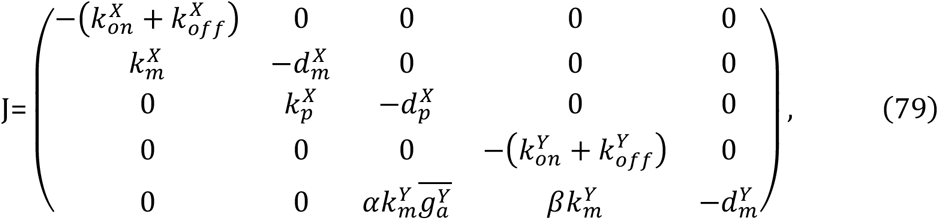

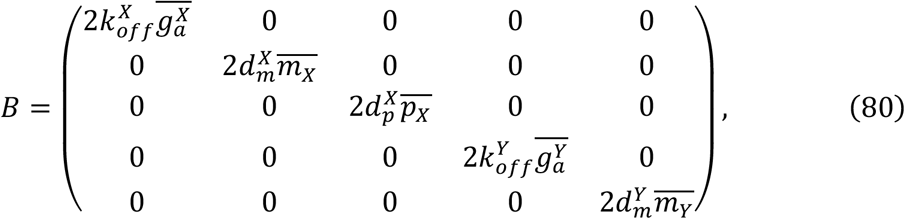

where 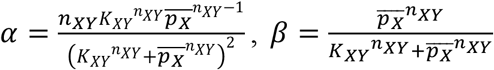, and 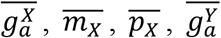, and 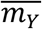 were obtained by solving stationary equations in the **Model Description** above. Then, the covariance matrix, *C*, was obtained by solving the matrix equation (Eq. 78) with elements *C_ij_* with indices *i* and *j* (1, 2, 3, 4, and 5 corresponding to the variables, 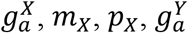, and *m_Y_*):

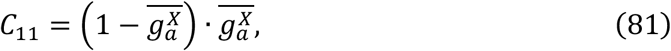

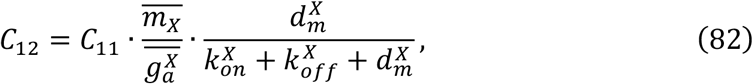

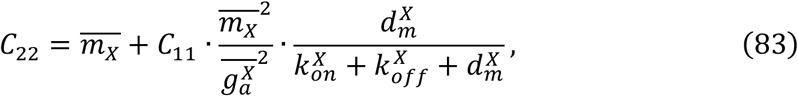

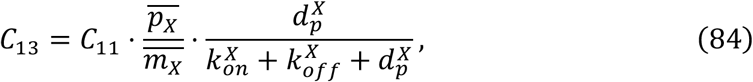

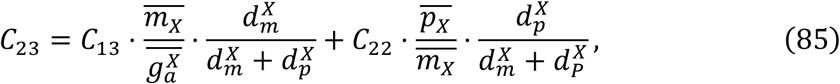

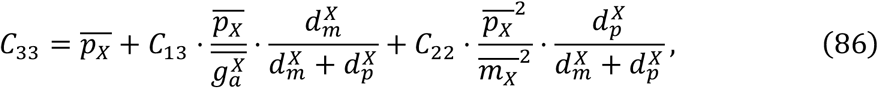

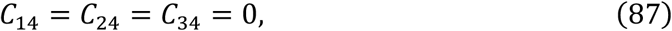

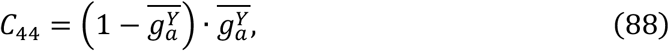

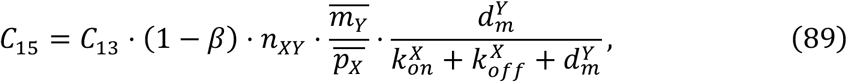

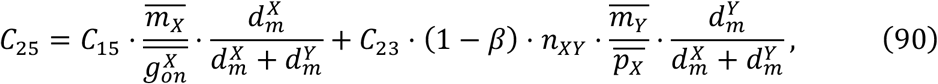

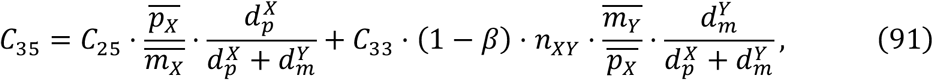

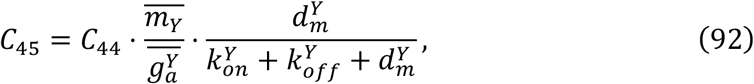

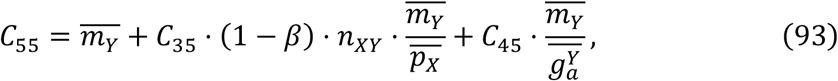

Finally, the analytical approximation of *Corr*(*m_X_,m_Y_*) was obtained as:

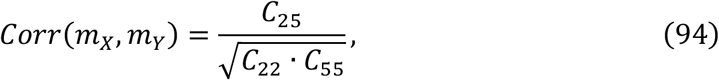

where *C*_22_ and *C*_55_ are the variances of mRNA X and mRNA Y, respectively.

Further expansion of the covariance between mRNAs X and Y, *C*_25_, revealed:

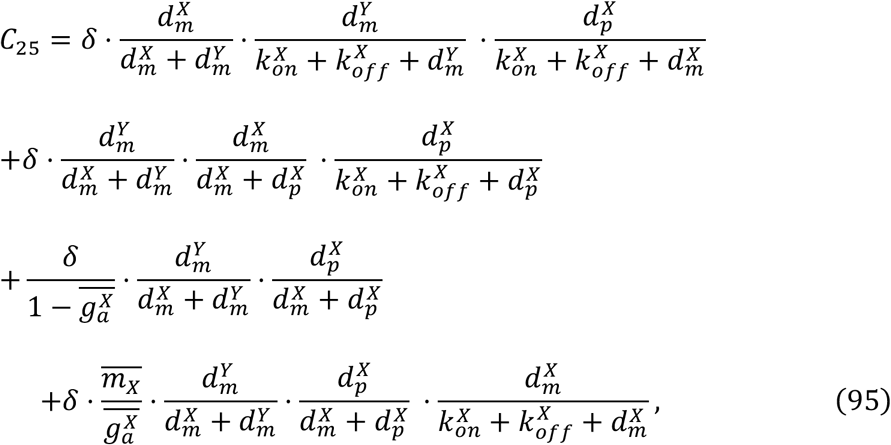

where 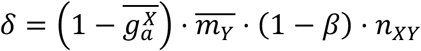, showing 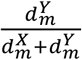 as one of the main contributors to *Corr*(*m_X_, m_Y_*) according to this analytical treatment.

### Deterministic modeling of the two-gene negative feedback circuit: Bifurcation analysis at a fixed point

We explored the phase space and bifurcation behaviors of the two-gene negative feedback circuit (Figure 4) (*29*). Our goal is to delineate, using deterministic modeling only based on the **Model Description** above, whether a given parameter combination would result in damped oscillation, limit cycle oscillation, or stable steady state. First, the fixed points (i.e., 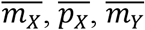, and 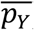) were obtained for each of parameter combinations by solving the stationary equations numerically shown in the **Model Description** above. Then, the Jacobian *J* at the fixed point was obtained as:

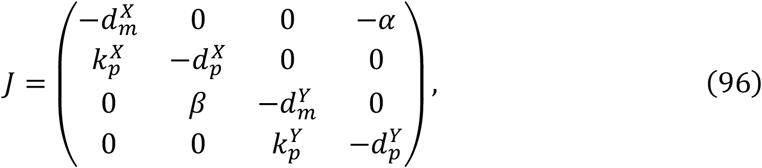

where

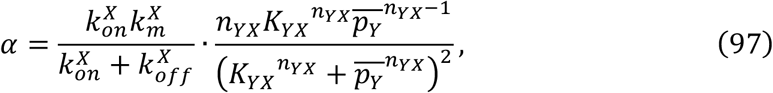

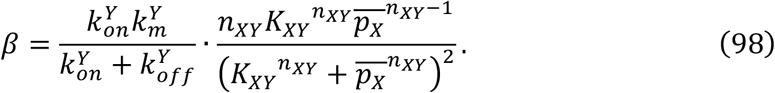

The eigenvalues *λ* of the Jacobian *J* at the fixed point were obtained by solving the characteristic equation of *J*:

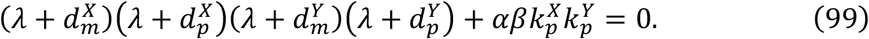

Depending on the parameter combination, this equation can have: 1) four real negative roots, 2) two real negative and two complex roots, or 3) four complex roots. Since the analytical approach for solving fourth degree polynomial equations is complex, we numerically solved this equation for each of the parameter combinations. More thorough treatment of four-dimensional Hopf bifurcation can be found in Asada and Yoshida (*59*).

The nature of the eigenvalues obtained for each parameter combination determine the behaviors of the circuit (*29, 60*). First, if the eigenvalues are all real and negative, then the system evolves to and settles at the stable fixed point (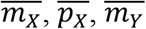, and 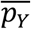). Second, if the eigenvalues include a pair or two pairs of complex values with negative real part, then, the system exhibits damped oscillation, eventually settling down to the stable fixed point. Lastly, if the eigenvalues include a pair of complex conjugates with positive real part, then the system exhibits limit cycle oscillations (Figures 4E and S6B). Note that it is impossible that the eigenvalues include two pairs of complex conjugates with positive real parts since the sum of all eigenvalues should be negative (i.e., 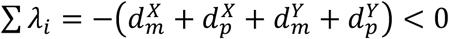).

### Power Spectral Analysis

The oscillatory behavior in the two-gene negative feedback network can be described in the frequency domain using power spectra analysis of the time trajectories. Power spectra were obtained by Fourier transformation of the auto-correlation functions or crosscorrelation functions as proven by the Wiener-Khinchin theorem (*15*). Ideally, we need a large number of realizations of the time trajectories to obtain the power spectra. However, here (and often in practice) we only have a single time trajectory and thus noise can be an issue. There are several methods to reduce such noise, including the averaging of multiple estimates from segments of the original trajectories and applying window functions for Fourier transformations (*61*). We employed both strategies together. We divided single time trajectories into multiple segments, applied Welch’s methods implemented in the *sapa* package in R to each of those segments, and averaged over multiple estimates. Based on the resolution needed and computation time, we set a limit on the length of the segment to not exceed 10^5^ time-points, which corresponds to ~8333 hours (10^5^ × 5 minutes) in the system’s time. Thus, this approach would miss anything that occurs in longer timescales or shorter than 5 minutes (the data acquisition time interval), which were not captured in the current power spectral analysis.

The unit of frequency in power spectra analysis is 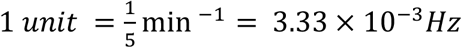 since the unit time interval is 5 min. The frequency range spanned several orders of magnitude. Thus, we applied the logarithm (with base 10) to define peak frequency (PF). PF spanned values ranging from −5 to −1, and the corresponding values of these in *Hz* spanned from 3.33 × 10^−8^ *Hz* to 3.33 × 10^−4^*Hz* using the general conversion formula; 10^PF^ × 3.33 × 10^−3^*Hz*. Intuitively, for example, if a periodic event occurs once every hour, the frequency is 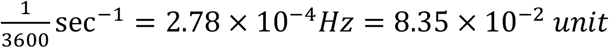, and after taking logarithm with base 10, PF = −1.08.

### The density color scale in scatterplots

The color scale (Figures 1F, 2A-C, 3B-C, S2D-E, S3A, S3C, S4A-D, S5B-C, S6A, and S6E-F) denotes the distribution density reflecting the relative abundance of data points. For each plot a 100 × 100 grid was constructed by partitioning the x and y ranges into 100 equal-size bins. The density value for each rectangle was obtained using the *kde2d* function in the R package *MASS* with the option n=100. Then, each data point was assigned the density value (and shown using the corresponding color scale) of the bin within which the data point is located.

**Table S1.** (Related to Figures 1, 3, and 4) Kinetic parameters with biologically plausible ranges from the literature.

| Parameter | Description (unit) | Range |
| --- | --- | --- |
| $k_{on}$ | The rate of switching of the promoter state from off- to on-state (event/hour) | 0.04-0.4 |
| $k_{off}$ | The rate of switching of the promoter state from on- to off-state (event/hour) | 0.01-0.5 |
| $k_m$ | The maximum rate of mRNA production (events/hour) | 0.1-3500<br>(0.1-1000)* |
| $d_m$ | The rate of mRNA degradation (events/molecule/hour) | 0.004-16 |
| $k_p$ | The rate of protein production (events/molecule/hour) | 0.1-20000<br>(0.1-1000)* |
| $d_p$ | The rate of protein degradation (events/molecule/hour) | 0.0005-6 |
| $n$ | Cooperativity for Hill functions of proteins (Hill coefficient) regulating the promoters/enhancers of downstream genes | 1<br>1-5<br>1, 5 |
| $K$ | The level (copy number or ratio to mean protein copy number) of the protein needed to achieve half maximum transcriptional rate for the downstream gene | 133.33 (copy number) for two-gene**<br>30-3000 (copy number) for two-gene negative feedback***<br>0.2-5 for (ratio) for three-gene feedforward |

Superscripts are used to denote the gene (for example, $k_{off}^X$ ) and subscripts are used to denote parameters involved in the interaction between genes, for example, $K_{XZ}$ and $n_{XZ}$ for interaction between protein X and the promoter/enhancer of gene Y).
\*For genes encoding transcription factors.
\*\*The value was fixed as our previous work (12).
\*\*\*In our analyses, these parameters were rescaled and expressed as a ratio to the estimated mean copy number of the protein. For further information including references, see STAR Methods.

## References

1. R. N. Germain, M. Meier-Schellersheim, A. Nita-Lazar, I. D. C. Fraser, Systems Biology in Immunology: A Computational Modeling Perspective. Annu. Rev. Immunol. 29, 527–585 (2011).

2. J. Gunawardena, Models in biology: ‘accurate descriptions of our pathetic thinking.’ BMC Biol. 12, 29 (2014).

3. A. C. Babtie, M. P. H. Stumpf, How to deal with parameters for whole-cell modelling. J. R. Soc. Interface. 14, 20170237 (2017).

4. R. N. Gutenkunst, J. J. Waterfall, F. P. Casey, K. S. Brown, C. R. Myers, J. P. Sethna, Universally Sloppy Parameter Sensitivities in Systems Biology Models. PLOS Comput. Biol. 3, e189 (2007).

5. F. Li, T. Long, Y. Lu, Q. Ouyang, C. Tang, The yeast cell-cycle network is robustly designed. Proc. Natl. Acad. Sci. 101, 4781–4786 (2004).

6. W. Bialek, Perspectives on theory at the interface of physics and biology. Rep. Prog. Phys. 81, 012601 (2018).

7. K. Baum, A. Z. Politi, B. Kofahl, R. Steuer, J. Wolf, Feedback, Mass Conservation and Reaction Kinetics Impact the Robustness of Cellular Oscillations. PLoS Comput. Biol. 12, e1005298 (2016).

8. H. Koeppl, M. Hafner, J. Lu, Mapping behavioral specifications to model parameters in synthetic biology. BMC Bioinformatics. 14, S9 (2013).

9. W. A. Lim, C. M. Lee, C. Tang, Design Principles of Regulatory Networks: Searching for the Molecular Algorithms of the Cell. Mol. Cell. 49, 202–212 (2013).

10. A. Tanay, A. Regev, Scaling single-cell genomics from phenomenology to mechanism. Nature. 541, 331–338 (2017).

11. M. B. Elowitz, A. J. Levine, E. D. Siggia, P. S. Swain, Stochastic Gene Expression in a Single Cell. Science. 297, 1183–1186 (2002).

12. A. J. Martins, M. Narayanan, T. Prüstel, B. Fixsen, K. Park, R. A. Gottschalk, Y. Lu, C. Andrews-Pfannkoch, W. W. Lau, K. V. Wendelsdorf, J. S. Tsang, Environment Tunes Propagation of Cell-to-Cell Variation in the Human Macrophage Gene Network. Cell Syst. 4, 379–392.e12 (2017).

13. A. Raj, A. van Oudenaarden, Stochastic gene expression and its consequences. Cell. 135, 216–226 (2008).

14. J. Elf, M. Ehrenberg, Fast Evaluation of Fluctuations in Biochemical Networks With the Linear Noise Approximation. Genome Res. 13, 2475–2484 (2003).

15. N. G. V. Kampen, Stochastic Processes in Physics and Chemistry, Third Edition (North Holland, Amsterdam; Boston, 3 edition., 2007).

16. D. T. Gillespie, Stochastic Simulation of Chemical Kinetics. Annu. Rev. Phys. Chem. 58, 35–55 (2007).

17. J. Paulsson, Summing up the noise in gene networks. Nature. 427, 415–418 (2004).

18. A. Raj, C. S. Peskin, D. Tranchina, D. Y. Vargas, S. Tyagi, Stochastic mRNA Synthesis in Mammalian Cells. PLoS Biol. 4, e309 (2006).

19. P. Thomas, N. Popovi, R. Grima, Phenotypic switching in gene regulatory networks. Proc. Natl. Acad. Sci. 111, 6994–6999 (2014).

20. R. Milo, S. Shen-Orr, S. Itzkovitz, N. Kashtan, D. Chklovskii, U. Alon, Network Motifs: Simple Building Blocks of Complex Networks. 298, 5 (2002).

21. S. Neph, A. B. Stergachis, A. Reynolds, R. Sandstrom, E. Borenstein, J. A. Stamatoyannopoulos, Circuitry and Dynamics of Human Transcription Factor Regulatory Networks. Cell. 150, 1274–1286 (2012).

22. S. Mangan, U. Alon, Structure and function of the feed-forward loop network motif. Proc. Natl. Acad. Sci. 100, 11980–11985 (2003).

23. L. Goentoro, O. Shoval, M. Kirschner, U. Alon, The incoherent feedforward loop can provide fold-change detection in gene regulation. Mol. Cell. 36, 894–899 (2009).

24. T. M. Filtz, W. K. Vogel, M. Leid, Regulation of transcription factor activity by interconnected, post-translational modifications. Trends Pharmacol. Sci. 35, 76–85 (2014).

25. U. Alon, Network motifs: theory and experimental approaches. Nat. Rev. Genet. 8, 450–461 (2007).

26. R. N. Germain, Maintaining system homeostasis: the third law of Newtonian immunology. Nat. Immunol. 13, 902–906 (2012).

27. A. Becskei, L. Serrano, Engineering stability in gene networks by autoregulation. Nature. 405, 590–593 (2000).

28. J. P. Pett, M. Kondoff, G. Bordyugov, A. Kramer, H. Herzel, Co-existing feedback loops generate tissue-specific circadian rhythms. Life Sci. Alliance. 1, e201800078 (2018).

29. S. H. Strogatz, Nonlinear Dynamics and Chaos: With Applications to Physics, Biology, Chemistry, and Engineering, Second Edition (Westview Press, Boulder, CO, 2 edition., 2015).

30. A. Woller, D. Gonze, T. Erneux, The Goodwin model revisited: Hopf bifurcation, limit-cycle, and periodic entrainment. Phys. Biol. 11, 045002 (2014).

31. D. B. Forger, C. S. Peskin, Stochastic simulation of the mammalian circadian clock. Proc. Natl. Acad. Sci. 102, 321–324 (2005).

32. H. Li, Z. Hou, H. Xin, Internal noise stochastic resonance for intracellular calcium oscillations in a cell system. Phys. Rev. E. 71 (2005), doi:10.1103/PhysRevE.71.061916.

33. A. J. McKane, J. D. Nagy, T. J. Newman, M. O. Stefanini, Amplified Biochemical Oscillations in Cellular Systems. J. Stat. Phys. 128, 165–191 (2007).

34. N. Guisoni, D. Monteoliva, L. Diambra, Promoters Architecture-Based Mechanism for Noise-Induced Oscillations in a Single-Gene Circuit. PLOS ONE. 11, e0151086 (2016).

35. D. Gonze, W. Abou-Jaoudé, The Goodwin Model: Behind the Hill Function. PLOS ONE. 8, e69573 (2013).

36. O. Shoval, H. Sheftel, G. Shinar, Y. Hart, O. Ramote, A. Mayo, E. Dekel, K. Kavanagh, U. Alon, Evolutionary Trade-Offs, Pareto Optimality, and the Geometry of Phenotype Space. Science. 336, 1157–1160 (2012).

37. J. A. J. Arpino, E. J. Hancock, J. Anderson, M. Barahona, G.-B. V. Stan, A. Papachristodoulou, K. Polizzi, Tuning the dials of Synthetic Biology. Microbiology. 159, 1236–1253 (2013).

38. D. G. Loyola R, M. Pedergnana, S. Gimeno García, Smart sampling and incremental function learning for very large high dimensional data. Neural Netw. 78, 75–87 (2016).

39. L. Breiman, Random Forests. Mach. Learn. 45, 5–32 (2001).

40. A. Verikas, A. Gelzinis, M. Bacauskiene, Mining data with random forests: A survey and results of new tests. Pattern Recognit. 44, 330–349 (2011).

41. P. Mohammadi, N. Beerenwinkel, Y. Benenson, Automated Design of Synthetic Cell Classifier Circuits Using a Two-Step Optimization Strategy. Cell Syst. 4, 207–218.e14 (2017).

42. L. Dolken, Z. Ruzsics, B. Radle, C. C. Friedel, R. Zimmer, J. Mages, R. Hoffmann, P. Dickinson, T. Forster, P. Ghazal, U. H. Koszinowski, High-resolution gene expression profiling for simultaneous kinetic parameter analysis of RNA synthesis and decay. RNA. 14, 1959–1972 (2008).

43. M. Jovanovic, M. S. Rooney, P. Mertins, D. Przybylski, N. Chevrier, R. Satija, E. H. Rodriguez, A. P. Fields, S. Schwartz, R. Raychowdhury, M. R. Mumbach, T. Eisenhaure, M. Rabani, D. Gennert, D. Lu, T. Delorey, J. S. Weissman, S. A. Carr, N. Hacohen, A. Regev, Dynamic profiling of the protein life cycle in response to pathogens. Science. 347, 1259038–1259038 (2015).

44. G.-W. Li, D. Burkhardt, C. Gross, J. S. Weissman, Quantifying Absolute Protein Synthesis Rates Reveals Principles Underlying Allocation of Cellular Resources. Cell. 157, 624–635 (2014).

45. L. Mariani, E. G. Schulz, M. H. Lexberg, C. Helmstetter, A. Radbruch, M. Löhning, T. Höfer, Short-term memory in gene induction reveals the regulatory principle behind stochastic IL-4 expression. Mol. Syst. Biol. 6 (2010), doi:10.1038/msb.2010.13.

46. M. Rabani, J. Z. Levin, L. Fan, X. Adiconis, R. Raychowdhury, M. Garber, A. Gnirke, C. Nusbaum, N. Hacohen, N. Friedman, I. Amit, A. Regev, Metabolic labeling of RNA uncovers principles of RNA production and degradation dynamics in mammalian cells. Nat. Biotechnol. 29, 436–442 (2011).

47. B. Schwanhäusser, D. Busse, N. Li, G. Dittmar, J. Schuchhardt, J. Wolf, W. Chen, M. Selbach, Global quantification of mammalian gene expression control. Nature. 473, 337–342 (2011).

48. R. Milo, R. Phillips, Cell Biology by the Numbers (Garland Science, New York, NY, 1 edition., 2015).

49. I. M. Sobol’, On the distribution of points in a cube and the approximate evaluation of integrals. USSR Comput. Math. Math. Phys. 7, 86–112 (1967).

50. M. D. Mckay, R. J. Beckman, W. J. Conover, A Comparison of Three Methods for Selecting Values of Input Variables in the Analysis of Output from a Computer Code. Technometrics. 42, 55 (2000).

51. T. Homma, A. Saltelli, Importance measures in global sensitivity analysis of nonlinear models. Reliab. Eng. Syst. Saf. 52, 1–17 (1996).

52. A. Saltelli, M. Ratto, S. Tarantola, F. Campolongo, Update 1 of: Sensitivity Analysis for Chemical Models. Chem. Rev. 112, PR1–PR21 (2012).

53. D. T. Gillespie, The chemical Langevin equation. J. Chem. Phys. 113, 297–306 (2000).

54. D. Kwiatkowski, P. Phillips, P. Schmidt, Y. Shin, Testing the null hypothesis of stationarity against the alternative of a unit root: How sure are we that economic time series have a unit root? J. Econom. 54, 159–178 (1992).

55. L. Breiman, Bagging Predictors. Mach. Learn. 24, 123–140 (1996).

56. G. Zhang, Y. Lu, Bias-corrected random forests in regression. J. Appl. Stat. 39, 151–160 (2012).

57. A. Liaw, M. Wiener, Classification and regression by randomForest. R News. 2, 18–22 (2002).

58. G. J. Berman, D. M. Choi, W. Bialek, J. W. Shaevitz, Mapping the stereotyped behaviour of freely moving fruit flies. J. R. Soc. Interface. 11, 20140672 (2014).

59. T. Asada, H. Yoshida, Coefficient criterion for four-dimensional Hopf bifurcations: a complete mathematical characterization and applications to economic dynamics. Chaos Solitons Fractals. 18, 525–536 (2003).

60. Y. Kuznetsov, Elements of Applied Bifurcation Theory (Springer, New York, 3rd edition., 2004).

61. D. B. Percival, A. T. Walden, Spectral Analysis for Physical Applications (Cambridge University Press, 1 edition., 1993).

